# Cell cycle difference creates cortical tension difference that separates germ layer fates

**DOI:** 10.1101/2021.11.10.468043

**Authors:** Naohito Takatori, Yuuya Tachiki

## Abstract

The segregation of germ layer fates is a fundamental step for embryogenesis, but the underlying molecular mechanism is unclear. In ascidians, mRNA localization coupled to nuclear migration and subsequent asymmetrical partitioning of the mRNA separates mesodermal and endodermal fates. The lack of quantitative characterization of nuclear and mRNA localization has hindered our understanding of the molecular basis of fate separation. Here, we quantitatively examined the movement of the nucleus and changes in cell shape and found that the nucleus moves to the mesodermal cell side across the region of the future cleavage furrow. However, this migration was not decisive for the asymmetric distribution of *Not* mRNA. Asymmetry of surface tension, caused by cell cycle difference between animal and vegetal hemisphere cells, deformed the mesendoderm cell and determined the position of the cleavage furrow, thereby ensuring the asymmetric partitioning of *Not* mRNA and segregation of fates. This study demonstrates how cell cycle control and the physical force relationships between cells are involved in the segregation of developmental fates.

## Introduction

Segregation of germ layer fates into distinct cells establishes the foundation for further specification of cell fates in descendant cells. The mesoderm and endoderm fate separation mechanism has been studied extensively at the single-cell level in *Caenorhabditis elegans* [1, 2]. Recent studies in an ascidian, *Halocynthia roretzi*, also addressed how the two germ layer fates are separated at the single-cell level [3-5]. Lineage-specific expression of *Lhx3* and *ZicN*, each required and sufficient for specification of endoderm or mesoderm fates[3, 4, 6-8], respectively, is regulated by a homeodomain transcription factor, Notochord (Not), whose transcript localized to the future mesoderm region of the mesendoderm cell and was subsequently partitioned to the mesoderm daughter (Figure 1A). Localization of *Not* mRNA was coupled to the nuclear migration that was directed toward the mesoderm side by PI3Kα signaling [5]. However, the lack of quantitative data on nuclear migration and *Not* mRNA localization has hindered further understanding of the processes and mechanisms underlying nuclear migration toward the mesoderm side.

**Figure 1.**
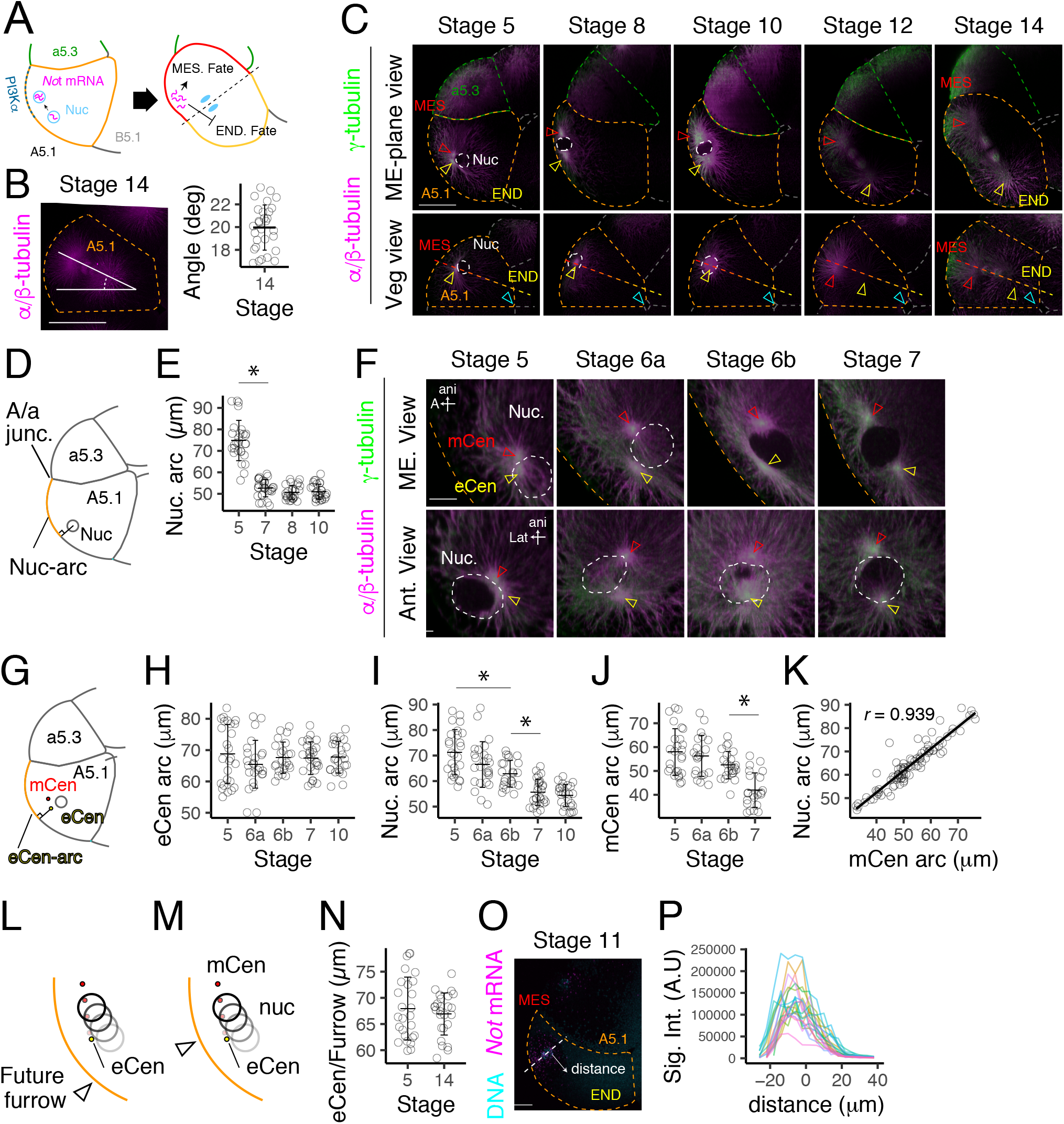
The mesendoderm cell nucleus moves toward the mesoderm side. (A) Schematic diagram of mesoderm and endoderm fate separation in ascidian embryo. (B) Spindle orientation at Stage 14, when the furrow is formed. The mesendoderm cell is indicated by orange dots. Anterior to the left. Vegetal view. (C) Nucleus and centrosome positions during the 16-cell stage. Lateral and vegetal view of the same cell are shown. The nucleus is indicated by white dots. A5.1 and a5.3 cells are outlined by orange and green dots, respectively. In the lateral view, nucleus and centrosomes were observed in the ME-plane. The relative position of the graded dots and cyan arrowhead (vegetal pole) are the same throughout. Anterior to the left and animal pole to the top in lateral view, Anterior to the left and medial side of the embryo to the bottom in vegetal view. Images that show the eCen and mCen are overlayed in vegetal view images at stages 12 and 14. (D) Diagram showing how nucleus position was defined. (E) Length of the Nuc-arc. (F) Close-up of the nucleus and centrosomes during nuclear migration. Lateral view and vegetal view of the same embryos are shown. Animal pole to the top. (G) Diagram showing how the eCen position was defined. (H-J) Length of the eCen-arc, Nuc-arc and mCen-arc from stage 5 to stage 10. (K) Correlation between Nuc- and mCen-arc from stage 5 to stage 7. (L, M) Diagram showing how the movement of the nucleus may not contribute to positioning the nucleus in the future mesoderm-side to the cell. (N) Position of the eCen (stage 5) and cleavage furrow (stage 14). Mesendoderm cell and neighbor a5.3-cell are marked by orange and green dots, respectively. White broken lines show outline of the nucleus. Red and yellow hollow arrowheads indicate mCen and eCen, respectively. Cyan arrowheads indicate the vegetal pole: intersection of A5.1/A5.1 contact area and A5.1/B5.1 contact area. MES: mesoderm side of the mesendoderm cell, END: endoderm side. Scale bar = 50 µm (B, C); 10 µm (F, O). Error bars represent one standard deviation.

Blastomeres display significant changes in morphology during embryogenesis. Recent studies have reported the role of cell morphology in defining mitotic spindle orientation and position in ascidian embryos [9, 10]. Surface tension has been shown to be a significant cause of such change, and its roles in forming the morphology of the embryo have been investigated [11-14]. The role of these morphological changes and cortical tension differences among cells in fate separation has begun to be understood [15]. In this work, we present results demonstrating that nuclear movement of the mesendoderm cell indeed places the nucleus and *Not* mRNA in the future mesoderm side of the cell. Subsequently, the mesendoderm cell underwent a change in shape that was critical in determining the furrow position. The difference in cell-cycle length caused this deformation via a difference in cortical tension between the mesendoderm cell and its neighbor cell. Our results suggest a profound relationship between cell cycle-dependent asymmetry in cortical tension that determines the cleavage furrow and nuclear migration position, which together contributes to fate separation.

## Results

### The mesendoderm cell divides along the axis that deviates 20 degrees from the embryo’s medial plane

The first goal of the study was to reevaluate in 3D space whether the mesendoderm cell nucleus moves toward the mesoderm pole. The direction of the mesoderm-endoderm axis was examined by observing the orientation of the mitotic apparatus. The mitotic apparatus was oriented 20 degrees relative to the medial plane of the embryo at Stage 14 (Figure 1B), when a cleavage furrow began to be formed (Movie S1F, Figure 1C). Since the direction of the spindle determines how cellular components are partitioned between mesoderm and endoderm cells, we decided to reevaluate the asymmetry within the cell along this axis.

### The nucleus appears to move toward the mesoderm side along the mesoderm-endoderm axis

We examined the asymmetry of nucleus position along the mesoderm-endoderm axis in 3D images of fixed mesendoderm cells from stage 0 to stage 14 (Movie S1B-S1F). Preliminary analysis suggested that nuclear migration and spindle formation occurs in a single plane that is tilted 20 degrees relative to the medial plane of the embryo, which is herein called the “ME-plane” (Figure S1A). Time-lapse observation revealed that when ME-plane is defined as a plane that includes the endoderm-side centrosome (eCen) at stage 5, the nucleus is observed in the ME-plane at stages 5 to 11. We found that eCen can be observed by MitoTracker staining (Figure S1C). eCen signal was detected in the ME-plane from stages 5 to 8 (Figure S1B). We confirmed the position of the eCen in fixed embryos by immunostaining with anti-γ-tubulin antibody. The planes that includes the eCen or the spindle were at the same relative position to the posterior-medial end of the cell (vegetal view Figure 1C, cyan arrowhead: posterior-medial end of the cell). These results support that the eCen and spindle are in the same plane, the ME-plane, from stages 5 to 14. When nuclear position was examined in the ME-plane at stage 5, the two centrosomes, the mesoderm-side centrosome (mCen) and eCen, were on the cell-surface-side of the nucleus (Figure 1C, Movie S1C). By stage 8, the nucleus was near the cell surface, and the centrosomes were on both sides of the nucleus (Figure 1C, Movie S1D). The mesendoderm cell underwent a change in shape around stages 8 and 10 (Figure 1C, Movie S1E). The nucleus was observed up to stage 10. At stage 11, the mesendoderm cell entered the M-phase (Figure S1D). The mitotic spindle was observed at stage 12, and the cleavage furrow was formed at stage 14 (Figure 1C, Movie S1F).

### The A5.1/a5.3-cell junction can be used as a reference point to measure the position of the nucleus

We examined whether we could use the A5.1/a5.3-cell junction (A/a junction, Figure 1D) to quantify the position of the nucleus. Carbon particles placed at the A5.1/a5.3-cell junction remained in the junction from stage 5 to 14 and were detected at the junction of the two cells under a confocal microscope (Figure S1E, white arrowhead). The position of the nucleus was defined using the length of the arc in the ME-plane (Figure 1D). The position of the nucleus was examined up to stage 10 because A5.1 cells enter M-phase at stage 11 (Figure S1D). The nucleus moved toward the mesoderm side from stage 5 to 7 (Table 1, Figure 1E).

### The nucleus moves to the mesoderm side of the eCen

At stage 5, the two centrosomes adjoined at the medial side of the nucleus (Figure 1F, Movie S1C) but parted from stage 5 to 7 (Table 2, Figure 1F, S1F, Movie S1C, S1D, S1G), and straddled the nucleus at stage 7. Statistical analysis did not support the movement of eCen toward the mesoderm side (Figure 1G, 1H, S1G, Table 3). eCen also did not move toward the medial plane of the embryo (Table 4, Figure S1F). The nucleus moved toward the mesoderm side (Table 5, Figure 1I) and the medial side (Table 6, Figure S1H). mCen moved toward the mesoderm side from stage 5 to stage 7 (Table 7, Figure 1J). A strong correlation was observed between the movement of the nucleus and that of mCen (Figure 1K). Together these results suggest that the nucleus, along with mCen, moves to the mesoderm side of eCen.

### The nuclear movement positions the nucleus in the future mesoderm region

The nuclear movement shown above may not be important for localizing *Not* mRNA in the future mesoderm region: if the centrosomes and the nucleus are already in the future-mesoderm region at stage 5, the movement may not be essential for *Not* mRNA localization (Figure 1L). On the other hand, if the nucleus and centrosomes are in the future-endoderm region at stage5, the movement could be important (Figure 1M). Statistical analysis did not support the difference between the position of the eCen at stage 5 and the furrow at stage 14 (Figure 1N, *p* = 0.466; *g* = -0.201 (0.345, -0.757)). We labeled the endoderm side of the nucleus at stage 7, and the labels were detected near the centrosome (Figure S1I, S1J, S1K). When labeled embryos were cultured to stage 15, particles were in the cleavage furrow. These results suggest that the position of the eCen at stage 5 is in the vicinity of the furrow at stage 14 and that nuclear movement from stage 5 to 7 places the nucleus in the future mesoderm.

### Nuclear migration localizes the bulk of *Not* mRNA to the mesoderm daughter

Distribution of *Not* mRNA was examined at stage 11. Since the cell enters M-phase, and transcription would end at this stage, stage 11 is suited to examine the effect of nuclear migration on *Not* mRNA localization. *Not* mRNA was detected by whole-mount *in situ* hybridization (Figure 1O). Signal intensity and distance from the nucleus along the mesoderm-endoderm axis was measured. The mesoderm-endoderm axis was tentatively set to 45 degrees. The distribution of *Not* mRNA was skewed toward the mesoderm-side and most of the RNA signal was observed within 20 microns of the nucleus (Figure 1P).

### The mesendoderm cell A5.1 cell does not increase in volume or total surface area during nuclear migration and cell shape change

Our results thus far suggest that nuclear migration is essential for asymmetric partitioning of *Not* mRNA. We next examined how the cleavage furrow is positioned in relation to the nucleus. Previous studies in ascidians demonstrated the relationship between cell shape and the orientation/ position of the mitotic spindle at the 32-cell stage [12, 13]. We found that the mesendoderm cell underwent a significant change in cell shape (Figure 1C) and hypothesized that the change in A5.1 cell shape is involved in positioning the furrow. We first examined the nature of the change in cell shape and then suppressed the change to examine its role in furrow positioning. We noticed that the surface area of the A5.1 cell facing the perivitelline space might increase from stages 5 to 10.

Growth in cell size or membrane synthesis may cause an increase in the surface area facing the perivitelline space. We measured the volume and surface area of the A5.1 cell from the 3D images, but the statistical analysis did not support differences after stage 5 (Figure 2A, 2B, Table 8, 9). The cell surface area facing the perivitelline space decreased from stage 3 to 5 and increased from stage 8 to 12 (Table 10, Figure 2C). Contrarily, the contact area between A5.1 cell and B5.1 cell (Figure 2D, 2E) increased from stage 3 to 5 and then decreased from stage 10 (Table 11, Figure 2F). These results are consistent with the shape change caused by compaction and decompaction rather than cell growth or membrane synthesis. The contact area between A5.1 and a5.3 cells (Figure 2D) did not change from stage 5 to 14 (Table 12, 13, Figure 2G, S2A: two different data sets). The surface area at stage 10 may be different from that at stage 12, but statistical analyses are not conclusive. These results are consistent with our results showing that labels placed at A5.1/a5.3 cell junction remain in the junction from stage 5 to stage 14.

**Figure 2.**
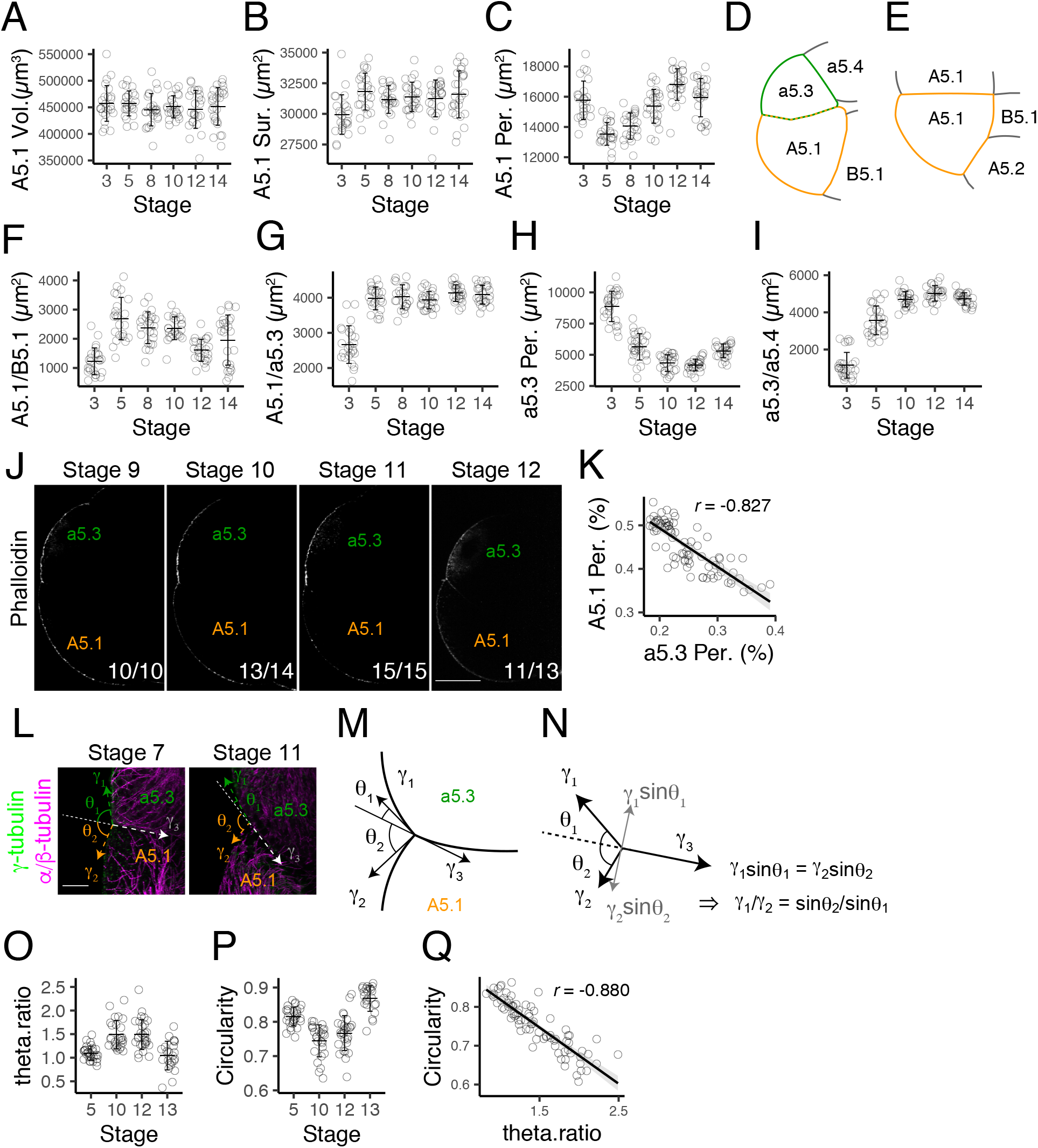
The A5.1-cell is deformed by the a5.3-cell during de-compaction. (A, B) A5.1 cell volume and total cell surface area from Stage 3 to Stage 14 measured from 3D images. (C) Surface area of the A5.1 cell facing the perivitelline space. (D, E) Spacial relationship between 16-cell stage blastomeres. (D): ME-plane view, (E): vegetal view. (F, G) Surface area of the A5.1 cell facing the B5.1 cell and a5.3 cell. (H) Surface area of the a5.3 cell facing the perivitelline space. (I) Surface area of the a5.3 cell facing the a5.4 cell.(J) Surface actin cytoskeleton stained with phalloidin at the 16-cell stage. (K) Relationship between surface areas of A5.1 and a5.3 cells. (L, M) Contact angles at A5.1/a5.3 cell junction. (N) Diagram explaining the equation for calculating surface tension ratio. (O) Surface tension ratio during the 16-cell stage. (P) Circularity of the A5.1 cell section at the ME-plane. (Q) Correlation between surface tension ratio and circularity in the A5.1. Scale bar = 50 µm (J); 10 µm (L). Error bars indicate one standard deviation.

### The cell shape change is accompanied by the convergence of the cell’s long axis

We examined the distribution of the cell long axis to characterize the change in cell shape (Figure S2B). Points that form the long axes (see Materials and Methods for details) were dispersed in the posterior and anterior regions of the cell at the early 16-cell stage but converged after stage 10. The angle of the long axis relative to the medial plane of the embryo was also initially spread but converged after stage 10 and coincided with that of the spindle at stage 14 (Figure S2B, Rose diagram). The lateral angle of the long axis also gradually converged around stage 10 (Figure S2C). These results suggest that the change in cell shape leads to convergence of the cell’s long axis.

### The neighbor a5.3 cell also undergoes compaction and decompaction, but at a different pace

Analysis of the volume and the surface area of the a5.3 cell supported that cell size does not change (Table 14, 15, Figure S2D, S2E). The cell surface area facing the perivitelline space decreased from stage 3 to 10 and then increased from stage 12 to 14 (Table 16, Figure 2H). The contact area between a5.3 and a5.4 cells (Figure 2D) increased from stage 3 to 12 and then decreased till stage 14 (Table 17, Figure 2I). These results show that the a5.3 cell undergoes compaction and decompaction at a slower schedule than the A5.1 cell.

### The difference in decompaction timing is accompanied by the surface tension difference between A5.1 and a5.3 cells

Previous studies showed that surface actin is critical for surface tension which plays a central role during compaction [16]. Studies using ascidian showed that surface tension differences deform vegetal hemisphere cells at the 32-cell stage [13]. We examined the relationship between a5.3 and A5.1 cells because the contact area between A5.1 and a5.4 cells is around one-tenth or less of the contact area between A5.1 and a5.3 (Figure S2F). We found that the surface actin signal is stronger in the a5.3 cell than the A5.1 at stages 10 and 11 (Figure 2J). The ratio of surface tension was estimated using contact angles at the A5.1/a5.3 cell junction (see Materials and Methods, Figure 2L, 2M, 2N). The tension ratio (a5.3/A5.1) increased from stage 5 to 10 and decreased from stage 12 to 13 (Table 18, Figure 1O). The circularity of the section of the A5.1 cell decreased from stage 5 to 10 and increased from stage 12 to 13 (Table 19, Figure 2P) and showed a strong correlation with the estimated tension ratio (Figure 2Q) supporting that tension difference causes cell deformation in A5.1 cells.

### The surface tension difference between a5.3 and A5.1 cells deforms the A5.1 cell

We tested whether difference in surface tension causes the deformation of the A5.1 cell by ectopic expression of a dominant-negative form of Rho protein (T19N-Rho) in the a5.3 cell. RNA encoding T19N-Rho was injected into both a5.3 and a<u>5.3</u>-cells around stage 2 to stage 4 (Figure 3A). Surface actin signal decreased in the injected cells (Figure 3B), and surface area facing the perivitelline space increased in injected cells (Figure 3C: *p* = 0.00115; *g* = 0.983 (1.63, 0.446)). This was not accompanied by the increase in cell volume (Figure S3A: *p* = 0.32; *g* = -0.249 (0.264, -0.787)) or total surface area (Figure S3B: *p* = 0.999; *g* = 0.000256 (0.526, -0.502)). Consistent with the change in compaction of the a5.3 cell, the a5.3/a5.4 cell surface are decreased (Figure S3C: *p* = 7.57e-07; *g* = -1.62 (−1.09 ,-2.36)). Surface tension ratio between a5.3 and A5.1 cells decreased (Figure 3D: *p* = 0.00172; *g* = -1.22 (−0.872, -1.9)).

**Figure 3.**
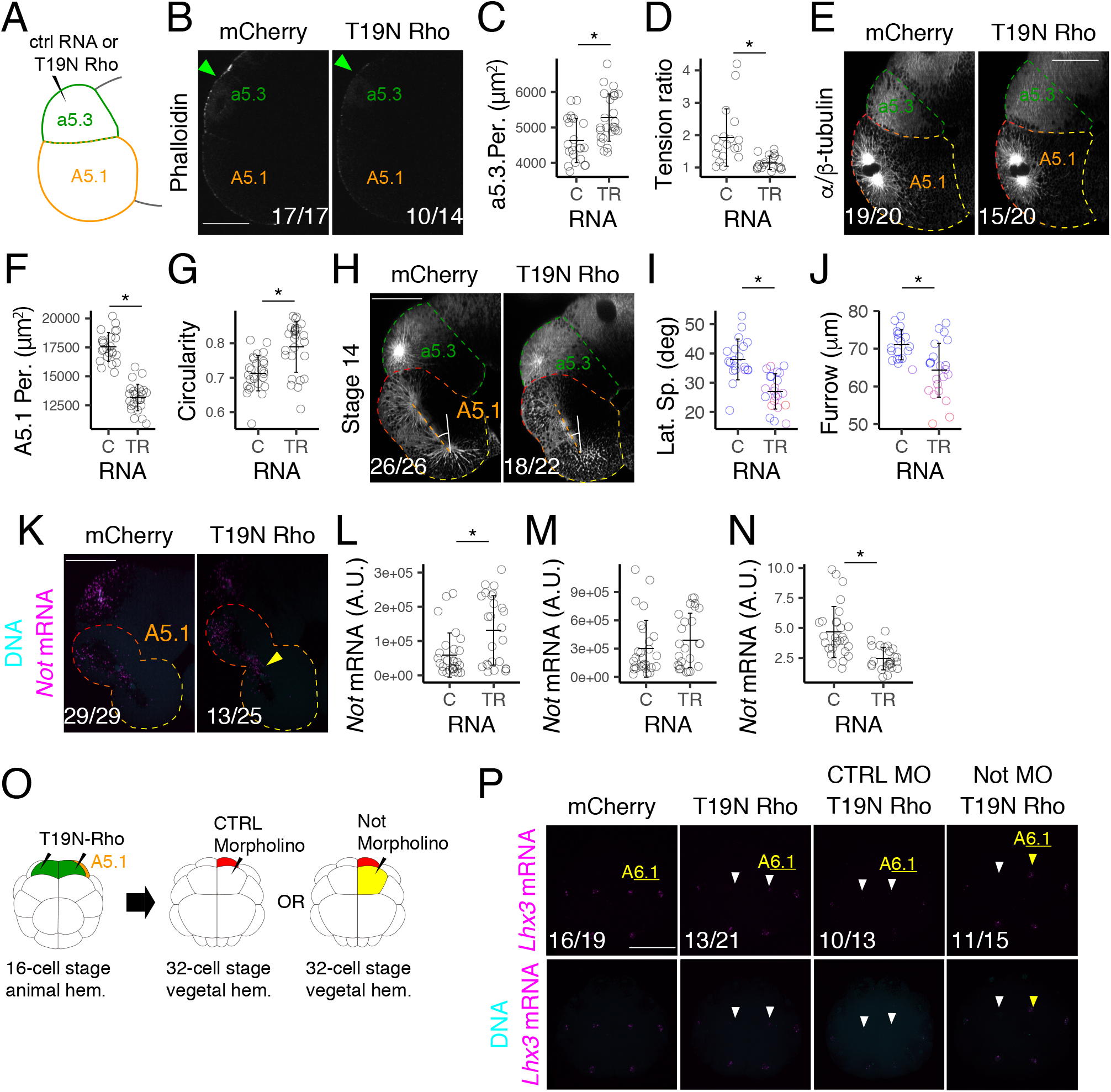
a5.3 cell surface tension deforms the A5.1-cell and separates fates. (A) Schematic diagram showing how T19N-Rho was injected into the a5.3 cell. (B) Surface actin cytoskeleton stained by phalloidin in injected embryos. (C) Surface area of the a5.3 cell facing the perivitelline space. (D) Surface tension ratio between A5.1 and a5.3 cells in injected embryos. (E) Representative example of morphology of A5.1 cells in injected embryos at Stage 10. Lateral view, anterior to the left and animal pole to the top. (F) Surface area of the A5.1 cell facing the perivitelline space. (G) Circularity of the A5.1 cell section at the ME-plane. (H) Morphology and the orientation of the spindle in A5.1 cells. Embryos were immunostained with anti-tubulin antibody. The angle between the spindle and the A5.1/B5.1 junction is shown. (I) Angle between the spindle and the A5.1/B5.1 contact surface. (J) The distance between the cleavage furrow and the A5.1/a5.3 cell junction on the cell surface. Color is defined by position of the furrow. (K) Distribution of *Not* mRNA at Stage 15. Yellow arrowhead indicates *Not* mRNA in the future endoderm region. Future mesoderm and endoderm cells are indicated by red and yellow dots, respectively. (L) Sum of *Not* mRNA signal intensity in the future endoderm region of the cell at Stage 15. (M) Sum of *Not* mRNA signal intensity in the A5.1 cell at Stage 15. (N) Diagram showing the experiment scheme. T19N-Rho was injected into a5.3 and a<u>5.3</u>-cells at the 16-cell stage. Then, control or Not MO was injected into the A<u>6.1</u> cell at the 24-cell stage. (O) Expression of *Lhx3* at the 32-cell stage. White arrowheads indicate loss of *Lhx3* expression in endoderm lineage cells. Yellow arrowheads indicate *Lhx3* mRNA detected in cells injected with Not MO. C: Control RNA; TR: T19-Rho RNA. Per: Perivitelline space. Spi.: spindle. MO: Morpholino Oligo. A.U.: Arbitrary Unit. Numbers in the panels indicate the proportion of cells that are represented by the image. Scale bar = 50 µm (B, D, H, K, P). Error bars indicate one standard deviation.

We examined the shape of the A5.1 cell at stage 10 because it is the final stage when the asymmetry of the nucleus, which translates to the asymmetry of *Not* mRNA localization, can be observed. The shape of the A5.1 cell at stage 10 differed in T19N-Rho injected embryos (Figure 3E, S3D). The surface area facing the perivitelline space decreased (Figure 3F: *p* = 0; *g* = -3.62 (−2.94, -4.78)), and the circularity of the section in the ME-plane increased (Figure 3G: *p* = 7.58e-05; *g* = 1.19 (2.05, 0.615)), quantitatively confirming that the shape has changed. The cell long axes did not converge in T19N Rho RNA injected embryos (Figure S3D). These changes was not accompanied by increased or decreased cell volume (Figure S3E: *p* = 0.394; *g* = -0.21 (0.294 ,-0.714)) or total surface area (Figure S3F: *p* = 0.626; *g* = -0.12 (0.357 ,-0.642)).

### Surface tension difference between a5.3 and A5.1 cells orients mitotic spindles and determines furrow position

To test the notion that surface tension difference determines the furrow position in the A5.1 cell, we examined the orientation of the spindle and furrow position at stage 14. The shape of the A5.1 cell at stage 14 was altered in T19N Rho injected embryos (Figure 3H, S3G). The cell long axes did not converge in some T19N Rho RNA injected embryos, and the spindle orientation was aberrant (Figure S3G, S3H, middle). In other embryos, the cell long axis did converge but at a different angle, and the spindle also formed at a similar angle (Figure S3G, S3H, right). The angle of the spindle to the medial plane in all injected embryos the angles became divergent in T19N-Rho injected embryos (F test, *p* = 4.60e-06), but the difference in the mean of the angles were not supported by statistical analysis (Figure S3I: *p* = 0.791; *g* = 0.0779 (0.641, -0.518)). On the other hand, the anterior tilt of the spindle in the ME-plane decreased (Figure 3H, 3I: *p* = 3.86e-06; *g* = -1.65 (−1.04, -2.5)). We found that cleavage furrow position was shifted toward the mesoderm side in T19N-Rho injected embryos (Figure 3J: *p* = 0.000534; *g* = -1.13 (−0.596, -1.89)). In some embryos, the furrow position was in the region where the nucleus would be at stages 7 to 10 (Figure 1E, 3J).

### A parsimonious model explains the furrow position using lateral spindle angle, vegetal spindle angle, and surface tension

To understand how the position of the furrow is determined, we examined the relationship between lateral angle, vegetal angle, and furrow position. We found that the furrow position shifts toward the endoderm side when the vegetal angle approaches 20 degrees or when the lateral angle increases (Figure S3K, S3L). No clear relationship was found between the vegetal and lateral angles (Figure S3M). Models that explain the relationship between the angles of the spindle and the furrow position were explored using AIC criteria (Table 20). We found that the inclusion of surface tension as an explanatory factor decreased AIC (models a vs. b and c vs. d). The square term of the vegetal angle also explained the data better than the vegetal angle (models b vs. d), which is visible in the relationship between the vegetal angle and the furrow (Figure S3K, S3M). Adding interaction term between lateral angle and the square of the vegetal term to the model did not improve the AIC value (Table 20). Thus, the model

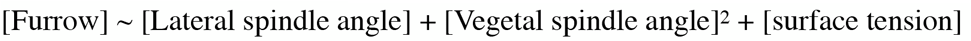

explains the data best (Table 20). The regression plane of the formula is shown in Figure S3K and shows that the furrow shifts toward the endoderm side as the vegetal angle approaches 20 degrees and the lateral angle increases.

### The difference in surface tension between a5.3 and A5.1 cells contributes to *Not* mRNA partitioning

*Not* mRNA was detected by whole-mount *in situ* hybridization, and the signal intensity was measured in the future-mesoderm and endoderm side of the cell (Figure 3K). The sum of signal intensity in the future-endoderm cell increased (Figure 3L: *p* = 0.00409; *g* = 0.846 (1.51, 0.315)) without the change in total signal intensity (Figure 3M: *p* = 0.281; *g* = 0.294 (0.885, -0.225)) in T19N-Rho RNA injected embryos. Mis-partitioning of *Not* mRNA, however, could be caused by disruption of nuclear migration up to stage 7. Statistical analysis did not support the difference in nucleus position in T19N-Rho and control RNA injected embryos (Figure S3O: *p* = 0.743; *g* = 0.0855 (0.624, -0.43)), supporting that mis-partitioning was caused by diminished deformation of A5.1 cell.

### The difference in surface tension contributes to fate separation

Ectopic expression of Not in endoderm cells leads to loss of expression of Lhx3, a transcription factor required and sufficient for endoderm fate, in the endoderm A6.1 cell [4, 5]. We tested whether cortical tension difference is involved in fate specification in the A6.1 cell. First, we determined the timing within the 32-cell stage to examine *Lhx3* expression. Gene expression can be easily mischaracterized at the 32-cell stage due to differences in the timing of entrance to the M-phase among cells. mRNA of many genes are detected in the nucleus in ascidians. Therefore, signals appear weaker in cells that have entered the M-phase due to loss of nuclear membrane and dispersion of mRNA. We examined the timing of phosphorylation of histone H3 in the 32-cell stage embryo and found that the 70 min embryo is consistently not immunostained with an anti-pH3 antibody (Figure S3Q). Thus, we decided to examine the expression of *Lhx3* at 70 min. Embryos injected with T19N-Rho showed loss of *Lhx3* mRNA signals in A6.1 cells (Figure 3P). Loss of *Lhx3* expression was rescued by injection of Not MO into the A6.1-cell, supporting the notion that loss of *Lhx3* expression is due to mis-partitioned *Not* mRNA (Figure 3P).

### Cell cycle progression difference between a5.3 and A5.1 cells deforms the A5.1 cell

Asynchrony in the cell cycle creates cortical tension differences between animal and vegetal blastomeres and is crucial for mitotic spindle alignment at the 32-cell stage [12, 13]. We found that the compaction and decompaction timing of a5.3 and A5.1 cells are different (Figure 2) and hypothesized that the cell cycle asynchrony is the cause of cortical tension difference and determines the furrow position (Figure 4A). We first began by characterizing the difference in cell cycle progression between a5.3 and A5.1-cells at the 16-cell stage. Both a4.2 and A4.1 cells, mother cells of a5.3 and A5.1 cells, respectively, formed the cleavage furrow at stage 1 (Figure S4A).

**Figure 4.**
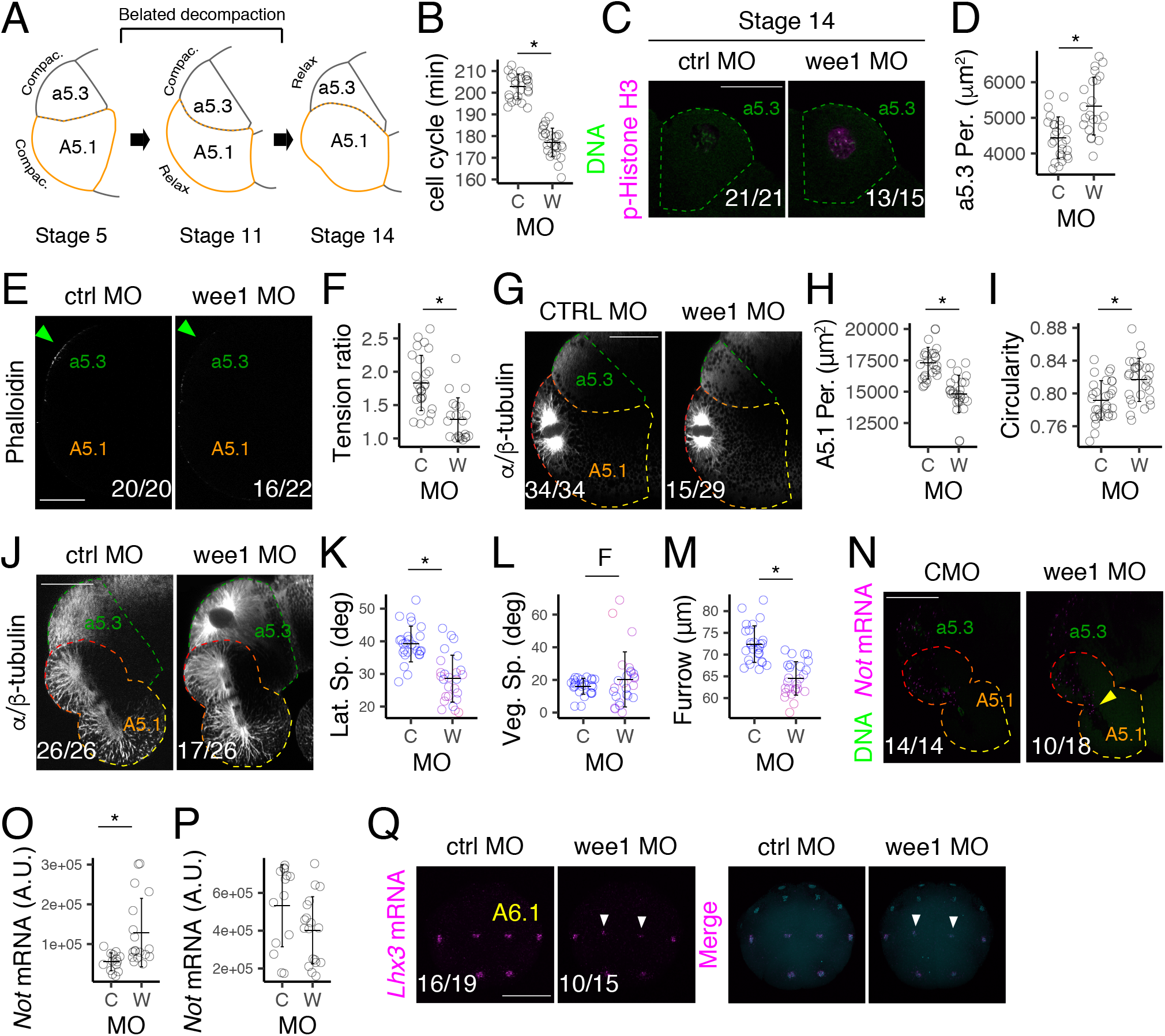
Cell-cycle difference separates mesoderm and endoderm fates. (A) Schematic diagram showing the model of A5.1 cell-deformation during the 16-cell stage. (B) Length of cell cycle in MO injected embryos. Formation of the furrow was observed under a stereomicroscope. (C) phosphorylation of Histone H3 in a5.3 cells at Stage 14 in MO injected embryos. Anterior to the front. Medial to the left. (D) Surface area of the a5.3 cell facing the perivitelline space. (E) Surface actin cytoskeleton (green arrowhead) at stage 10. (F) Surface tension ratio during the 16-cell stage. (G) Morphology of the A5.1 cell in injected embryos at Stage 10. (H) Surface area of the A5.1 cell facing the perivitelline space. (I) Circularity of the A5.1 cell section at the ME-plane. (J) Lateral view of the A5.1 cell in injected embryos at Stage 14. The angle between the spindle and the A5.1/B5.1 contact area is shown. (K) Angle between the spindle and the A5.1/B5.1 cell contact surface. Color is defined by position of the furrow show in (M). (L) Angle between the spindle and the medial plane: the A5.1/A5.1 cell contact surface. (M) The position of the cleavage furrow measured from the A5.1/a5.3 cell junction.(N) Distribution of *Not* mRNA at Stage 15. Yellow arrowhead indicates *Not* mRNA in the future endoderm region. (O) Sum of *Not* mRNA signal in the future endoderm region of the cell at Stage 15. (P) Sum of *Not* mRNA signal intensity in both mesoderm and endoderm region at Stage 15. (Q) Expression of *Lhx3* mRNA at the 32-cell stage. White arrowheads indicate loss of *Lhx3* expression in endoderm lineage cells. (E, F, J, N) Anterior to the left. Animal pole to the top. (C, Q) Vegetal view. Anterior to the top. C: Control MO; W: wee1 MO. MO: Morpholino Oligo. Per: Perivitelline space. Spi.: spindle. A.U.: Arbitrary Unit. F: F-test.. Numbers in the panels indicate the proportion of cells that are represented by the image. Scale bar = 50 µm. Error bars indicate one standard deviation.

Centrosome duplication occurred at stage 3 in both a5.3 and A5.1 cells (Figure S4B). Towards the end of the cell cycle, Histone H3 phosphorylation was detected at stages 16 and 11 in a5.3 and A5.1 cells, respectively (Figure S4C). Cleavage furrow formed at stage 20 and stage 14 in a5.3 and A5.1 cells, respectively (Figure S4D). Thus, the cell cycle progression is different not because the cell cycle is shifted later in the animal hemisphere but because the cell cycle is longer in the animal hemisphere (Figure S4E). Previously, knockdown of wee1 was used to resolve the asynchrony in another ascidian, *Phallusia mammillata* [12]. Injection of wee1 MO into both a5.3 and a<u>5.3</u>-cells resulted in a decrease, but not loss, of cell cycle progression difference in *H. roretzi* (Figure 4B). Histone H3 phosphorylation was detected prematurely in the a5.3 cell at stage 14 in wee1 MO-injected embryos (Figure 4C).

### Cell cycle asynchrony deforms vegetal hemisphere cells at the 16-cell stage

Surface area of the a5.3 cell facing the perivitelline space increased in injected cells (Figure 4D: *p* = 0.00888; *g* = 0.75 (0.267, 1.29)) but was not accompanied by increase in cell volume or total surface area (Figure S4F, *p* = 0.32; *g* = -0.249 (0.264 ,-0.787), Figure S4G, *p* = 0.999; *g* = 0.000256 (0.526, -0.502)). Cortical actin signal decreased in the injected cells (green arrowhead, Figure 4E). The surface tension ratio between a5.3 and A5.1 cells decreased (Figure 4F: *p* = 7.14e-06; *g* = -1.43 (−0.887, -2.2)), and A5.1 cell shape changed less in wee1 MO-injected embryos than controls at stage 10 (Figure 4G, S4J). Surface area of the A5.1 cell facing the perivitelline space decreased (Figure 4H: *p* = 4.98e-09; *g* = -1.93 (−2.67, -1.46)), and the circularity of the section of the A5.1 cell increased (Figure 4I: *p* = 0.00164; *g* = 0.875 (0.41, 1.49)). The cell long axes did not converge in wee1 MO-injected embryos (Figure S4J, Rose diagram). These changes were not accompanied by increase or decrease in cell volume (Figure S4H: *p* = 0.937; *g* = 0.0233 (−0.504, 0.652)) or total surface area (Figure S4I: *p* = 0.398; *g* = 0.236 (−0.332, 0.765)), supporting that A5.1 cell deformation is caused by change in the a5.3 cell cycle.

### Cell cycle progression difference orients mitotic spindles and determines furrow position at the 16-cell stage

Cleavage furrow was formed in the A5.1 cell at stage 14 in wee1 injected embryos, suggesting that cell cycle of the A5.1 cell is not affected (Figure 4J). Following the decrease in cell deformation at stage 10, the anterior tilt of mitotic spindles decreased in wee1 MO-injected embryos at stage 14 (Figure 4J, 4K: *p* = 6.06e-08; *g* = -1.74 (−2.65, -1.12)). When viewed from the vegetal pole, the cell long axes converged at aberrant angles in wee1 MO-injected embryos, and the spindles were oriented in similar directions (S4K, S4L). When the orientation of the spindle was examined in all injected embryos, spindles were less aligned in wee1 MO-injected embryos (Figure 4L: F test, *p* = 3.015e-08), but the difference in average angles was not supported by statistical analysis (Figure 4L: *p* = 0.234; *g* = -0.332 (0.248, -0.735)). The position of the furrow at stage 14 shifted toward the mesoderm side in MO-injected embryos (Figure 4J, 4M: *p* = 5.04e-09; *g* = 1.93 (2.61, 1.45)). We hypothesized that the difference in furrow position could be caused by difference in spindle length, but difference in spindle length was not supported by statistical analysis (Figure S4M: *p* = 0.787; *g* = -0.0742 (0.475, -0.601)).

### A parsimonious model that explains the furrow position using spindle angles and cell cycle differences

Similar to results in T19N-Rho injected embryos, furrow position shifted toward the endoderm side when the vegetal angle approached 20 degrees or when the lateral angle increased (Figure S4N, S4O). No clear relationship was found between the vegetal and lateral angles (Figure S4P). When models were selected using the AIC criteria, the inclusion of cell cycle difference as an explanatory factor decreased the AIC (Table 21), and the use of the square term of the vegetal angle instead of the vegetal angle consistently decreased the AIC. Since the orientation of the spindle in a 3D cell could be influenced by the shape of the cell, we included the interaction term between lateral and vegetal angles, but this did not decrease the AIC. Therefore, the formula : [Furrow] ∼ [Lateral spindle angle] + [Vegetal spindle angle]^2^ + [cell cycle difference] is the model with the smallest AIC (Table 21). The regression plane of the formula is shown in Figure S4P.

### Cell cycle asynchrony between a5.3 and A5.1 cells contributes to *Not* mRNA partitioning and fate separation

The sum of *Not* mRNA signal intensity in the future-endoderm cell increased in MO-injected embryos (Figure 4N, 4O: *p* = 0.00238; *g* = 1.08 (1.6, 0.75)), but analyses were inconclusive regarding the decrease in the total signal intensity (Figure 4P: *p* = 0.0819; *g* = -0.646 (0.0561, -1.56)). Embryos injected with wee1 MO showed loss or lessened *Lhx3* mRNA signals in A6.1 cells (Figure 4Q).

### Cortical tension acts downstream of the cell cycle asynchrony in changing A5.1 cell shape

Our results thus far suggest that cell cycle asynchrony between a5.3 and A5.1 cells causes cortical tension differences that result in cell shape change and correct positioning of the furrow in the A5.1 cell. We attempted to confirm this cascade by increasing cortical tension in wee1 MO-injected cells by injecting RNA encoding a constitutive active form of Rho (Q63L-Rho). Injection of Q63L-Rho RNA in the a5.3-cell resulted in excessive deformation of the A5.1 cell toward the animal pole (RNA: 20 pg/cell, Figure S5A). We adjusted the amount of Q63L-Rho RNA to 5-10 pg per cell and injected the RNA into the a5.3-cell after the injection of wee1 MO. The surface area of the a5.3-cell increased in wee1 MO-injected cells but decreased in wee1 MO and Q63L-Rho injected embryos (Table 22, Figure 5A). The surface tension ratio between a5.3 and A5.1 cells decreased with knockdown of wee1 but increased in Q63L-Rho co-injected embryos (Table 23, Figure 5C). Change in cell shape decreased in wee1 MO-injected cells but was similar to that of control embryos in Q63L-Rho co-injected embryos (Figure 5C). Cell long axes did not converge in wee1 MO-injected embryos but converged in embryos co-injected with Q63L Rho (Figure S5B, S5C). The surface area of the A5.1 cell facing the perivitelline space was rescued in Q63L-Rho injected embryos (Table 24, Figure 5D). The circularity of the section of the A5.1 cell also was rescued (Table 25, Figure 5E). None of these changes were accompanied by increased or decreased cell volume or total surface area in either a5.3 or A5.1 cells (Table 26, 27, 28, 29, Figure S5D, S5E, S5F, S5G).

**Figure 5.**
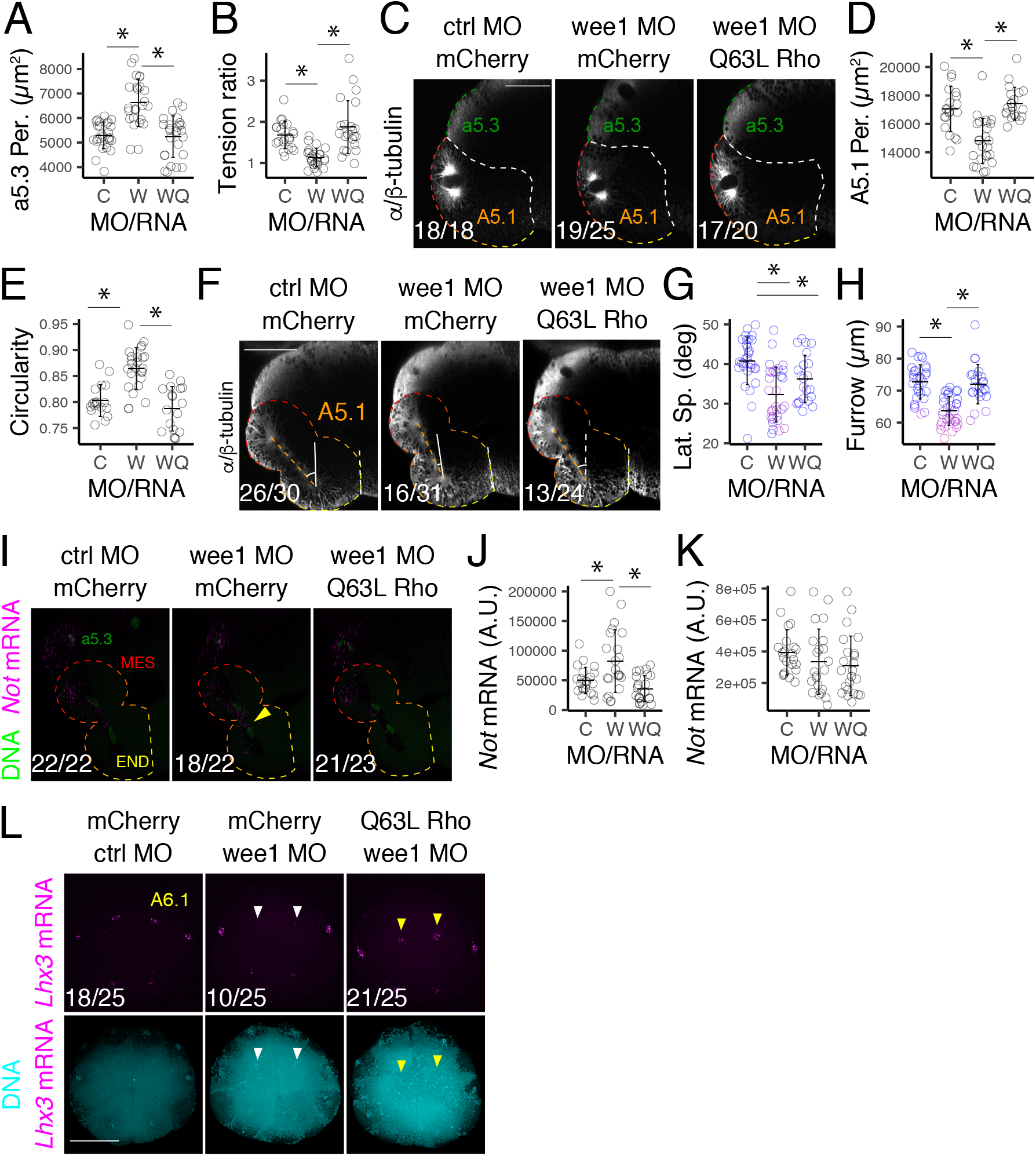
Cell-cycle difference separates fates by creating cortical tension difference. (A) Surface area of the a5.3 cell facing the perivitelline space. (B) Surface tension ratio in injected embryos. (C) Lateral view of the A5.1 cell in injected embryos at Stage 10. (D) Surface area of the A5.1 cell facing the perivitelline space. (E) Circularity of the A5.1 cell section at the ME-plane. (F) Lateral view of the A5.1 cell in injected embryos at Stage 14. Embryos were immunostained with anti-tubulin antibody. The angle between the spindle and the A5.1/B5.1 contact area is shown. (G) Angle between the spindle and the A5.1/B5.1 cell contact area. Color is defined by position of the furrow show in (H). (H) The position of the cleavage furrow. (I) Distribution of *Not* mRNA at Stage 15. Yellow arrowhead indicates ectopic *Not* mRNA in the future endoderm region. (J) Sum of *Not* mRNA signal intensity in the future endoderm region of the cell at Stage 15. (K) Sum of *Not* mRNA signal intensity in both the future endoderm and mesoderm regions at Stage 15. (L) Expression of *Lhx3* at the 32-cell stage. White arrowheads indicate loss of *Lhx3* expression in endoderm lineage cells. Yellow arrowheads indicate *Lhx3* mRNA detected in cells injected with Not MO. (A, F, I) Anterior to the left. Animal pole to the top. (L) Vegetal view. Anterior to the top. MO: Morpholino Oligo. Per: Perivitelline space. C: control MO and mCherry RNA; W: wee1 MO and mCherry RNA; WQ: wee1 MO and Q63L-Rho RNA. Spi.: spindle. MES: mesoderm; END: endoderm. Numbers in the panels indicate the proportion of cells that are represented by the image. Scale bar = 50 µm. Error bars indicate one standard deviation.

### Cortical tension acts downstream of the cell cycle asynchrony in determining furrow position and orienting the spindle

The anterior tilt of the spindle decreased in MO-injected embryos but recovered in Q63L-Rho RNA co-injected embryos (Figure 5F, 5G, Table 30). However, the difference between the mean of the spindle angle in control and the Q63L-Rho injected angle was supported by statistical analysis (Table 30). The shape of the cell viewed from the vegetal pole showed abnormality in wee1 MO-injected embryos, and the long axes were dispersed in several forms of disarray (two examples are shown in Figure S5H). The mean of the vegetal angle was not different in wee1 MO-injected embryos, but the difference in variance was supported by statistical analysis (Figure S5I, sd: 6.08 vs. 15.3, F-test: *p* = 3.025e-06). This was mitigated in Rho co-injected embryos, but statistical analyses are inconclusive (sd: 15.3 vs. 10.1, F-test: *p* = 0.0468). The position of the furrow at stage 14 shifted toward the mesoderm side in MO-injected embryos but shifted toward the endoderm side in Q63L-Rho co-injected embryos (Figure 5F, 5H, Table 31). Statistical analysis did not support the difference between control and Q63L-Rho co-injected embryos (Table 31).

### *Not* mRNA partitioning and fate separation are rescued in Q63L-Rho co-injected embryos

The signal intensity of *Not* mRNA in the future-endoderm region increased in MO-injected cells (Figure 5I, 5K, Table 32) but decreased in Q63L-Rho injected embryos. Differences in the whole-cell signal intensity were not supported by statistical analysis (Table 33, Figure 5L). Rescue of *Not* mRNA partitioning may have been because the nucleus migrated further to the mesoderm side in Q63L-Rho expressing embryos. However, the position of the nucleus did not change in wee1 MO or wee1 MO, and Q63L-Rho RNA injected embryos (Table 34, Figure S5J). Embryos injected with wee1 MO showed loss of *Lhx3* mRNA signals in A6.1 cells but expressed *Lhx3* in wee1 MO/Q63L-Rho RNA co-injected embryos. (Figure 5N).

## DISCUSSION

Our previous studies showed that localization and partitioning of *Not* mRNA, coupled with nuclear migration, separate germ layer fates but did not quantitatively establish that the nucleus moves toward the mesoderm side [4, 5]. Here, we showed that the nucleus indeed moves toward the future mesoderm side of the cell, but the movement is not decisive in itself and must later be accompanied by cell deformation that correctly positions the cleavage furrow.

Nuclear migration is often linked to asymmetric cell division that accompanies fate separation [17]. Our results demonstrate that the ∼ 15 µm nuclear movement from stage 5 to stage 7 moves the nucleus from the future-endoderm to the mesoderm region of the cell. Distribution of *Not* mRNA (Figure 1P) supports that the migration of the nucleus is sufficient to partition a large portion of *Not* mRNA to the mesoderm daughter cell. We also found that nuclear migration occurs near the region where the cleavage furrow is later formed (Figure 1N, S1P, S1Q, S1R), but the mechanism that brings the centrosome/nucleus to this region is not clear. There may be a structure near the future cleavage furrow, which attracts the centrosome and defines the cleavage furrow. In the posterior vegetal blastomeres, the centrosome-attracting body (CAB) attracts the centrosome and the nucleus, but similar structures are unknown in anterior cells, and we also did not identify any notable structure in this study. The localization of Kif2 and aPKC, which localizes to the CAB and is required to attract spindle poles in *Phallusia* [18], may be localized. Our observation suggests nuclear movement where microtubules contact the cortex and motor proteins pull the nucleus/ centrosome toward the cortex [17]. Cortical localization of dyneins and microtubule attachment factors Gai/LGN/NuMA complex [19], particularly that of NuMA [20], may determine the direction of eCen migration in ascidians.

Our results suggested that regions of eCen migration and furrow formation overlap, but the structure that attracts the centrosome if it exists, would have limited roles in defining the furrow position. Decreasing a5.3 cell surface tension shifted the furrow position (Figure 3I), but the nucleus position before cell deformation was not altered (Figure S3J, S3K). These results favor the notion that nuclear migration and furrow positioning are independent, but our results do not rule out the possibility that a single structure may loosely determine both the spindle and furrow positions. The mechanism defining the furrow position is further discussed below.

We found that the two centrosomes display different movements during nuclear positioning. mCen, but not eCen, migrated toward the mesoderm side with the nucleus, whose direction of migration is determined by PI3K signaling [5]. It is yet to be determined whether mCen movement is also directed by PI3K signaling. The difference in centrosome movements may reflect centrosome age [21, 22]. eCen consistently showed stronger γ-tubulin signals before nuclear migration at stages 3 to 5 (Figure 1C, 1F). Localization of age-specific proteins such as ninein and centrobin, which localize to mother and daughter centrosomes [21-23], respectively, may provide clues. Uncovering how the mCen and the nucleus move toward the future mesoderm side would be the central goal to understanding how mesoderm and endoderm polarity is formed in the mesendoderm cell. Interestingly, we noticed that the mesoderm-directed movement of the nucleus coincided with the separation of the centrosomes, and the nucleus protruded in between the separating nucleus during migration (Movies S1E). Centrosomes typically separate via sliding the two centrosomes along the nucleus surface [24], but in the *Halocynthia* mesendoderm cell, the relationship may be opposite, and centrosome separation mechanisms involving kinesin may play roles in nuclear migration.

Our results suggest that the deformation of the mesendoderm cell is caused by pulling force exerted on the mesendoderm cell from the neighboring a5.3 cell during decompaction. This confirms the conclusions of previous studies using *Phallusia* and *Ciona* [9, 10, 12, 13]. Using 3D images, we added that, at least in *Halocynthia* embryos, the cell volume and total surface area do not change during the deformation process (Figure 2), strengthening the view that the change in shape is a process that accompanies decompaction. Previous studies showed that the spindle direction is altered at the 32-cell stage [12]. We found that that the same holds at the 16-cell stage in *Halocynthia* embryos. Similar methods, expression of a dominant-negative form of Rho in a5.3-cells [13] or knockdown of wee1 [12, 13], were used in the current study and resulted in a change in cell morphology and spindle orientation at the 16-cell stage (Figure 3). These changes in cell morphology disturbed *Lhx3* expression in the endoderm cell lineage, supporting the role of cell deformation at the 16-cell stage in spindle positioning and cell fate determination. Discrepancy regarding the 16-cell stage in the current and previous work may be due to the difference in wee1 knockdown method, injection into the egg or to the a5.3-cell, or small sample size [12]. Also, the average spindle angle observed from the vegetal pole was also not different from that in controls (*p* = 0.234; *g* = -0.332 (0.251, -0.76)), and many embryos displayed spindle angles similar to those observed in the previous study [12].

Perturbation of the spindle angle at the 16-cell stage was likely caused by the change in cell shape in Rho or wee1 MO-injected embryos, as cell shape is a major factor in positioning the mitotic spindle [25, 26]. The cell long axis, which plays central roles in the shape-dependent orientation of the spindle, converged to the ME-plane during cell deformation and showed a strong correlation with the direction of the spindle (Figure S2). Dispersal of long axes that differ from each other by less than 1% of the total length seems to be a good indication that the cell lacks a clear unique long axis to which the spindle can align prior to stage 10. Mis-orientation of the spindle was coupled to a lack of convergence of cell long axes (Figure S3, S4), supporting that convergence of cell angle by cell deformation aligns the spindle. We found two patterns of cell long axis convergence, dispersed or converged to irregular angles, at at stage 14 (Figure S3G). The latter may have resulted from pulling by a5.4. The orientation of A5.1 cell spindle may be determined under balance between a5.3 and a5.4 cells. Our results support the notion that cortical tension difference aligns the spindle via deformation of the A5.1 cell and convergence of cell long axes.

Positioning of the cleavage furrow is determined by multiple factors, such as spindle orientation and polar relaxation [27-30]. We postulate the model of the positioning of the furrow as follows (Figure S6): the mesendoderm cell deforms as the result of higher cortical tension in the a5.3 cell (Stage 7-8 to 11), which brings the longest axis of the cell to 20 degrees from the medial plane of the embryo. The elongated shape of the cell enables the spindle to align to the cell’s long axis at stages 12-13. Equatorial stimulation from the spindle and polar relaxation concomitantly determine the furrow position. With wee1 knockdown, M-phase is initiated early, and cortical tension begins to relax in the a5.3 cell. A5.1 cell deformation is diminished, and the cell rounds up, resulting in misorientation of the spindle. The latter occurs with T19N-Rho injection as well. The furrow becomes misplaced by equatorial stimulation from the misaligned spindle and enhanced relaxation at the cortex (Figure S6). The relationship between spindle angles and furrow position was predicted by the same single model in both Figures 3 and 4. Interaction terms between cortical tension/cell cycle length and spindle angle did not improve the model (Table 20, 21), suggesting that cortical tension/cell cycle length difference does not affect the spindle angle’s effect on furrow position. The intercepts of cortical tension/cell cycle length were significantly different between control and treatment (intercept: -3.13, -5.18), suggesting that cortical tension and spindle angle independently regulate furrow position. The flow of cortical MyoII in the cortex has been shown to affect furrow position in *C. elegans* embryos [30, 31]. Cortical tension in a5.3 cells may exert force on the polar cortex of the A5.1 cell and thus, alter furrow position via a similar mechanism.

Our results point to the role of difference in the cell cycle between the a5.3 and A5.1 cells. However, it is not clear which phase of the cell cycle is longer in the a5.3 cell. Knockdown of wee1 hastened the cell cycle by about 20 minutes (Figure 4B, 4C, S4D) but did not abolish the cell cycle difference entirely as in *Phallusia* embryos [12]. The knockdown of wee1 may have been incomplete because wee1 MO was injected at the onset of the 16-cell stage. On the other hand, cell cycle length regulation may differ in *Halocynthia*, and length difference in G_1_ or S phase may account for the remaining difference. Cell cycle asynchrony at the 16-cell stage is a hallmark in a wide range of ascidian species [32, 33], but its cause is not clear. In *Halocynthia*, wee1 mRNA is distributed asymmetrically along the animal-vegetal axis [34], and this may be the cause of cell cycle length difference. However, it is not clear why the cell cycle lengths are not different at the 8-cell stage when the animal cell would have received more *wee1* mRNA compared to its vegetal sibling.

We found that cell deformation, *Not* mRNA partitioning, and fate separation could be rescued by increasing the tension in a5.3 cell in wee1 KD embryos (Figure 5, S5). These results suggest that cell cycle difference creates cortical tension difference that changes the A5.1 cell shape. However, the orientation of the mitotic apparatus was not fully rescued even when the furrow position was rescued (Figure 5I). These results may again reflect the contribution of polar cortex relaxation in determining the furrow position [27 -31].

In conclusion, we propose a model of mesoderm-endoderm fate separation in the *Halocynthia* embryo through quantitative analysis of cell morphology and nuclear position (Figure 6). The cell nucleus moves to the mesodermal cell side near the future cleavage furrow via centrosome movement. Nuclear migration is not solely decisive for the asymmetric partitioning of *Not* mRNA and requires cleavage furrow positioning through cell deformation. Cell deformation is caused by asymmetry of surface tension due to cell cycle differences between the animal and vegetal hemispheres and leads to the convergence of cell long axes. Spindle position is then determined by the cell long axis and cell shape. The position of the furrow is determined by the combined effects of spindle position and cortical tension. Our results illuminate the role of the cell cycle, cortical tension, and cell shape in the segregation of developmental fates.

**Figure 6.**
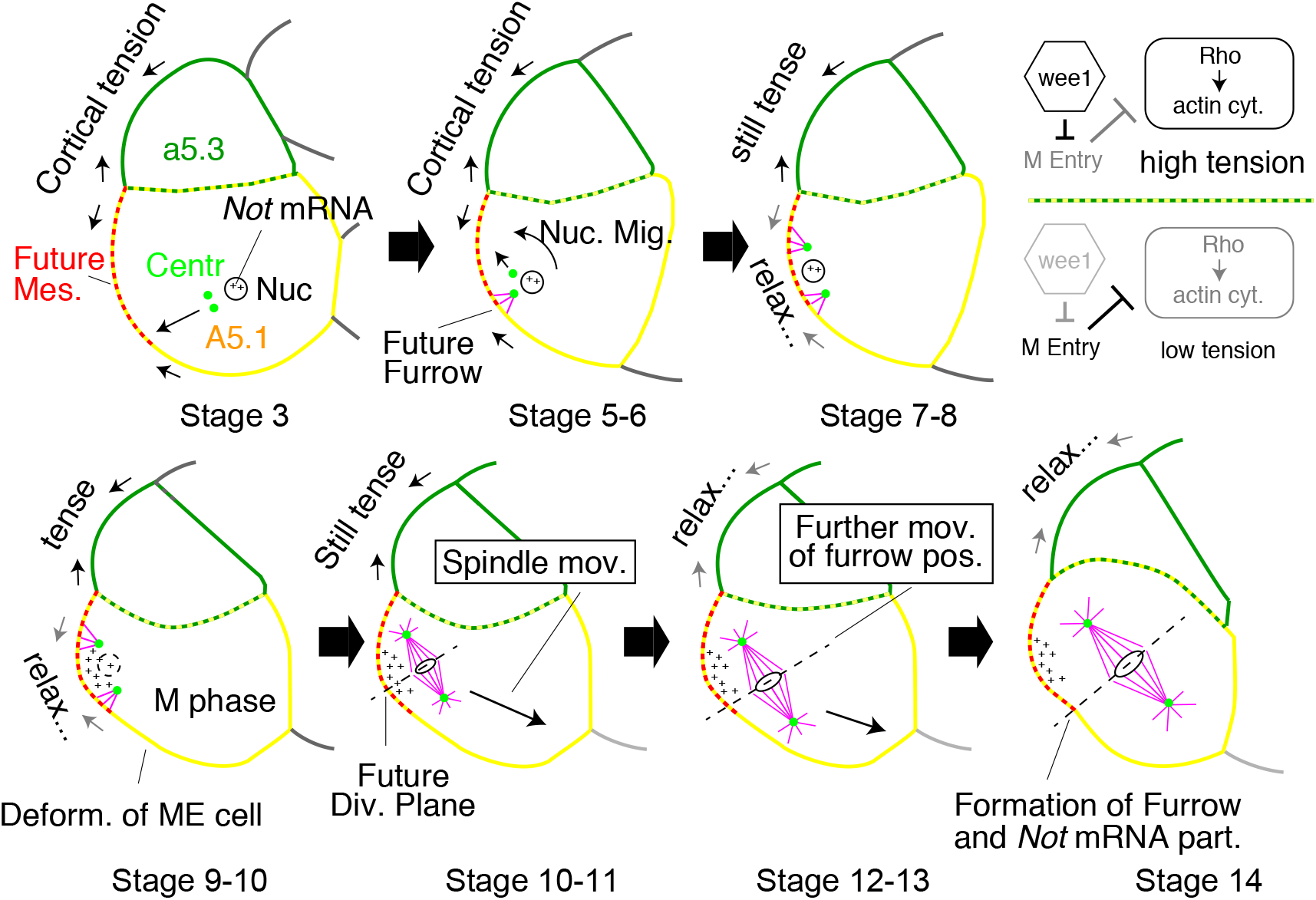
Model of fate separation by cell-cycle difference that creates cortical tension difference. Schematic model of *Not* mRNA partitioning via nuclear migration and cell deformation. Nuc: nucleus, Centr: centrosome, Mes: mesoderm, End: endoderm, ME: mesendoderm.

## Materials and methods

### Handling of animals and embryos

*Halocynthia roretzi* adults were purchased from fishermen in northern Japan, Aomori and Iwate prefectures in late November and December. Adult animals were kept in seawater at 7 degrees Celsius under constant light. Spawning was induced by raising the temperature to 11 degrees Celsius and exposing the animals to constant light after > 8 hr dark period. Spawned eggs were fertilized by artificial insemination. Embryos were kept at 7 degrees Celsius on gelatin coated glass bottom dishes (627861, Greiner Bio-One GmbH, Germany) and allowed to develop on temperature-controlled plates under a stereomicroscope equipped with 1X objective lens and 20X eyepiece (SMZ1270, NIKON, Japan). Embryonic development was synchronized by collecting animals that began to form the cleavage furrow in the B4.2 cell at the end of the 8-cell stage (Movie S1A). Embryos were devitellinated as described previously [4]. Stages represent minutes divided by 10 passed since the formation of the cleavage furrow (ex. 0 - 10 min: Stage 0, Supplemental Movie 1A).

### Immunohistochemistry

Embryos were fixed in 0.25% glutaraldehyde, 2% formaldehyde, 0.1 M PIPES pH 7.4, 50 mM EDTA pH 8.0, 300 - 350 mM sucrose, 10 mM MgSO_4_, 0.1 % Triton X-100 at 10 degrees Celsius for 12 to 16 hrs on incubator (WKN-9626, Waken, Japan) with gentle agitation. PIPES and EDTA were titrated with potassium hydroxide. Care was taken to introduce minimal sea water during fixation. Embryos were washed in PBS, 0.1% Triton X-100 (PBSTr), blocked in 0.5% blocking reagent/ PBSTr for 30 min in room temperature and incubated with primary antibodies for >16 h at 4 degrees Celsius. Embryos were washed in PBSTr and incubated in secondary antibodies: Alexa555 conjugated anti-rat antibody or MAX-PO(R) (Nichirei, Japan). Antibodies used were, YL1/2 (abcam, ab6160), anti-γ-tubulin antibody (ab179503), PH3 antibody (CST, 9701S), anti-Nup153 antibody (Covance, MMS-102P), anti-rat secondary antibody (ab150154), and MaxPO(R) (Nichirei, H1602). Embryos were counterstained using SYTOX green to visualize nuclear DNA as described previously [4].

### Phalloidin staining and whole mount *in situ* hybridization

Phalloidin staining and whole mount *in situ* hybridization were performed as described previously [4].

### Microscopy and image analysis

Embryos were mounted in 2’-2’ thiodiethanol [35] or methyl salicylate and placed on a custom made glass-bottom steel dish. Embryos were aligned so that the vegetal pole of the embryo faced the lens using human hair, which did not damage stained samples and was resistant to both mounting media. Images were obtained with 40 X Plan APO lambda or 60 X APO TIRF (NA 1.49) lens on a NIKON A1 confocal microscope (NIKON, Japan) mounted with piezo device (Physik Instrumente GmbH, Germany). Escape mechanism of the lenses were suppressed to prevent occasional erratic distortion of images along the z-axis. Cell dimensions were obtained by manual annotation of sections at each focal plane using custom scripts on ImageJ [36]. Coordinates of the centrosomes and spindle poles were manually obtained by identifying foci with highest γ-tubulin or α/β-tubulin signal. Coordinates of the nuclei in the ME-plane were obtained using the radius function in NIS-Elements (NIKON, Japan). Cell volume and surface areas of cells were calculated using custom R scripts [37]. Pairs of points that form the long axis of the cell were calculated using R from randomly chosen points on cell surface. 30,000 pairs were randomly chosen for calculation due to limitation in calculation capacity. Several rounds of calculations were performed to confirm results. Points that form the top fifty cell long axis were visualized by plotting onto 3D images using custom R and ImageJ scripts. Z-stack image and optical sections made from such images were shown in the Figures (ex. Figure S2B). The difference between the longest and shortest of the fifty axes were less than 1% of the length of the longest axis. Lateral and vegetal sections of A5.1 cell were obtained from 3D images using NIS-Elements. Quantification of mRNA signal was also performed using custom R and ImageJ scripts. Time-lapse observation of embryos were performed on NIKON A1 confocal microscope with CFI LWD 40 X WI lambda S lens. Sample temperature was controlled using stage top incubator KRiX-CSG (TOKAI-hit, Japan). Embryos were stained using MitoTracker ® Red CMXRos (100 nM in sea water, M7512, ThermoFisher) for time-lapse observation. Surface tension were estimated from contact angles at the a5.3/A5.1 celll junction was measured using Fiji software [38]. Modified Young-Dupre equation (γ_1_/γ_2_ = sinθ_2_/sinθ_1_) were used to determine the ratio of surface tension (Figure 2N).

### Injection of Morpholinooligo and RNA

Microinjection into unfertilized eggs and embryo were performed as described previously [4]. Injection into the 16- and 24-cell stage embryo was performed as follows: devitellinated embryos were placed in a well on gelatin coated plastic dish so the a5.3-cell or the A6.1 cell faced the lens. MO and RNA were injected into blastomeres using glass needles manipulated using MO-102 and MMO-220A (NARISHIGE, Japan). Injection was performed from stage 2 to stage 4. MO targeting Hr-wee1 (5’-AGCTTTAGCAGGACTGGTACATTGA-3’) was injected into both a5.3 cells (20 pg per cell). RNA was synthesized using mMESSAGE mMACHINE™ T3 Transcription Kit (AM1348, ThermoFisher) and also injected into both a5.3 cells. Amount of Rho RNAs were adjusted to 20 pg per cell. Sequence of MO against Not was reported previously [4]. Point mutation were induced in *Halocynthia* Rho (BAE95626) using PCR to create T19N and Q63L-Rho.

### Labeling of embryos

Embryos were labeled using carbon particles. First, carbon particles were placed in seawater, and particles of appropriate size were chosen under a stereomicroscope. Particles were then lifted and placed on embryos using human hair attached to a pipette tip. Embryos were allowed to sit for a few minutes to allow particles to attach to the cell surface. A5.1/a5.3 cell junction was labeled by orienting the embryo, so the junction faces the lens and placing the particle in the junction. A narrow groove of 100-micron depth was created on a gelatin-coated dish and used to orient the embryo. Labeling of the eCen region at stage 7 was performed as follows: embryos were placed, so the vegetal pole faces the lens. Carbon particles were placed adjacent to the caudal side of the nucleus under a stereomicroscope. A portion of the labeled embryos was fixed and observed under the confocal microscope. The remaining embryos were cultured to respective stages and observed under the stereomicroscope (Figure S1E, S1I) or confocal microscope (Figure S1E, S1J, S1K). Ten embryos were labeled in two different batches to confirm the results in both junction and eCen labeling.

### Statistical analysis

Statistical analyses were performed using R version 3.5.2 [37]. Sample sizes were determined using the power.t.test in pwr package: effect size, significance level and power were set at 0.8, 0.05 and 0.8, respectively. Welch and Games-Howell tests were used to compare means because these tests are known to be robust to non-normality, unequal sample size and unequal variances in underlying populations [39-41]. Combination of preliminary tests to determine the normality or variance and Student’s t-test was avoided on grounds of control over Type-I error [42]. Since normality and equal variance cannot always be assumed in cell biological observation, effect size was calculated to provide additional viewpoint on the difference between data [43]. Effect size, *g*, was calculated using Hedge’s method. 95% confidence interval of the effect size was calculated by bootstrapping (2000 re-iterations). 95% CI is shown after the effect size in parenthesis. Graphs were plotted using ggplot2 package. We searched for linear models that explain the mechanisms that determine furrow position using following explanatory variables: lateral angle of the spindle, vegetal angle of the spindle, surface tension of the a5.3 cell and cell cycle length of the a5.3 cell. Surface tension and cell cycle length were added as categorical variables. Lateral angle is the angle between the spindle and the animal-vegetal axis. Vegetal angle is the angle between the spindle and the medial plane. Squared terms of lateral and vegetal spindle angles and interaction effects between all variables were also included in the full model. Model selection was manually performed to identify a parsimonious model that explains the furrow position using the minimal AIC criteria [44, 45].

**Supplemental Figure 1: Supplementary figure to Figure 1**

(A) Schematic diagram showing the relationship between the ME-plane, centrosomes, nucleus and the mitotic spindle. Centrosomes are shown in yellow (eCen) or red (mCen). The nucleus and chromosomes are shown in green. The spindle is shown in magenta. The centrosome, nucleus and the spindle are in the ME-plane. (B) Snapshots from time-lapse 3D image of the 16-cell stage embryo stained with MitoTracker. Anterior to the top. Vegetal view. Mesendoderm cells are outlined with orange dots. Yellow arrowheads indicate eCen. We found that MitoTracker consistently stains centrosomes in *Halocynthia* mesendoderm cells (Figure S1C). ME-plane: graded dots from red to yellow and includes the eCen. The position of these dots are the same in all stages and indicate that the nucleus remains in the same optical plane (ME-plane) throughout these stages. (C) MitoTracker signals in the vicinity of the eCen. Representative images of the eCen in stage 5 embryos fixed and stained with MitoTracker and anti-α/β-tubulin antibody, respectively. (D) Stage 5 to 10 embryos stained with anti-α/ß-tubulin antibodies. Vegetal views of the same embryo at the focal plane that includes the nucleus (Nuc view) and the eCen (eCen view) at each stage. Anterior to the left and medial to the bottom. Cyan arrowheads indicate the posterior-medial end of the A5.1 cell is placed at the same place in each panels. (E) Overlay of stage 5 and 7, or stage 7 and 10, images that show the eCen. Images were aligned using the posterior-medial end of the A5.1 cell (cyan arrowhead). Note that the eCen remains in the ME-plane which is place in the same position in both panels. (F) Nuclei of the A5.1 cell at Stage 10 and 11. Embryos were immunostained with anti-p-Histone H3 and anti-Nup153 antibodies. White arrowheads indicate rupture in the nuclear membrane at Stage 11. (G) The angle of the A5.1/a5.3 cell junction relative to the horizontal plane (the plane that is perpendicular to the animal-vegetal axis). The junction was used to measure the positions of eCen and the nucleus. The angle indicates that error of 5 microns when placing the ME-plane to measure the nucleus position translates to 1.5 to 2 microns error in the nucleus position. (H) Distance between the eCen and the medial plane, A5.1/A5.1 cell contact surface. (I) The distance between eCen and mCen. (J) The distance between the nucleus and the medial plane of the embryo. (K) Representative image showing the results of labeling the A5.1/a5.3 cell junction with carbon particles. White arrowhead indicates the carbon particle placed at Stage 5. The particle remains in the junction at stage 14. The same particle was observed by confocal microscopy at Stage 14 and found to be at the junction of the A5.1/a5.3 cells. (L) Representative image showing the results of labeling the vicinity of eCen with carbon particles. Black arrowhead indicates the carbon particle placed at Stage 7. Black dots indicate the nucleus. The same embryo was observed at stage 15 (right side panel). Ten embryos were labeled. (M) A carbon particle was placed at the vicinity of the eCen at stage 7. Embryos were fixed and observed by confocal microscopy at Stage 7 and Stage 15. Embryos were stained with anti-α/ß-tubulin antibodies to visualize the eCen. Carbon particles are marked with cyan lines. Vegetal view. Anterior to the top and medial plane to the left. (N) 3D reconstruction of the image shown in (O). The carbon particle is near the eCen at Stage 7 and confirms the labeling of eCen. Particles are found in the cleavage furrow at Stage 15. Numbers in the panels indicate the proportion of cells that are represented by the image. Nuc: nucleus, MES: mesoderm, END: endoderm, ME: mesendoderm, eCen: endoderm-side centrosome, mCen: mesoderm-side centrosome. Scale bars = 10 µm (B, C, D, F, J) and 50 µm (E, K). Error bars in graphs indicate one standard deviation.

**Supplemental Figure 2 Supplementary figure to Figure 2**

(A) Surface area of the A5.1 facing the a5.3 cell. The data set is different to the set shown in Figure 2G. (B) Representative images showing the position of the pairs of points on the cell surface that forms the cell long axis. Top 50 long axis are shown using yellow (longest) to red (shortest) circles. Mesendoderm cells are outlined with white dots. Vegetal view. Anterior to the top, medial to the left. Rose diagrams showing the distribution of the longest axes in the representative embryo. Angle of the spindle in the representative embryo is shown in orange. (C) Lateral view of the same mesendoderm cells shown in (B). Points that were in the vicinity of the ME-plane are shown with orange dots. They are not necessarily from the pairs that form the long axis, so we show them differently to (B). Rose diagrams showing the distribution of the longest axis relative to the animal vegetal axis. (D) The volume of the a5.3 cell. (E) The total cell surface area of the a5.3 cell. (F) Relationship between contact area of a5.4 and A5.1 cells to contact are of a5.3 and A5.1 cells. Scale bars = 50 µm. Error bars indicate one standard deviation. (D) Surface area of the A5.1 facing the A5.2 cell.

(D) Surface area of the A5.1 facing the A<u>5.1</u> cell.

**Supplemental Figure 3 Supplementary figure to Figure 3**

(A) The cell volume of a5.3 cells in injected embryos at Stage 10. (B) The total surface area of a5.3 cells in injected embryos at Stage 10. (C) Surface area of the a5.3 cell facing the a5.4 cell. (D) Representative images showing the positions of the pairs of points on the cell surface that forms the cell long axis at stage 10. Top 50 long axis (∼0.1% of all calculated axes) are shown using yellow (longest) to red (shortest) circles. Mesendoderm cells are outlined with white dots. Vegetal view. Anterior to the top, medial to the left. Rose diagrams showing the distribution of the longest axes relative to the medial plane of the embryo. (E) The cell volume of A5.1 cells in injected embryos at Stage 10. (F) The total surface area of A5.1 cells in injected embryos at Stage 10. (G) The orientation of mitotic spindles at Stage 14 in injected embryos. Embryos were stained with anti-α/ß- tubulin antibodies to visualize the mitotic spindle. Representative image of the cell and positions of the pairs of points on the cell surface that forms the top 50 long axis. (H) Rose diagrams showing the distribution of the longest axes relative to the medial plane of the embryo. Angle of the spindle in the representative embryo is shown in orange. (I) The angle between the spindle and the medial plane viewed from the vegetal pole in all the injected embryos. (J) Rose diagrams showing the distribution of the longest axis relative to the animal vegetal axis in cells shown in Figure 3H. (K) Relationship between the position of the furrow to the angle between the spindle and the medial plane. (L) Relationship between the angle between the spindle and the A5.1/A5.1 contact area to the angle between the spindle and the medial plane. (M) Relationship between lateral and vegetal spindle angles. (N) Furrow position in response to the spindle angles. Red and blue surfaces correspond to T19N-Rho injected and control models, respectively. Surfaces represent the model prediction using the best AIC model. (O) The position of the nucleus at Stage 7. (P) Phosphorylation of histone H3 during the 32-cell stage. Time passed after the formation of the cleavage furrow in B5.2 is shown. Numbers in the panels indicate the proportion of cells that are represented by the image. Scale bar = 50 µm. Error bars indicate one standard deviation.

**Supplemental Figure 4 Supplementary figure to Figure 4**

(A) Formation of the cleavage furrow in A4.1 and a4.2 cells at Stage 1. Embryos were immunostained with anti-α/ß- and γ-tubulin antibodies. White arrowheads indicate cleavage furrows. (B) Centrosomes are duplicated at Stage 3 in both A5.1 and a5.3 cells. The orange hollow arrowheads indicate centrosomes before duplication. Red and yellow arrowheads indicate mCen and eCen, respectively. Green arrowheads indicate a5.3 cell centrosome. (C) Phosphorylation of Histone H3 at Stage 11 and Stage 16 in A5.1 and a5.3 cells, respectively. Embryos were immunostained with anti-pH3- and Nup153 antibodies. White arrowheads indicate pH3 signals. (D) Formation of the cleavage furrow in A5.1 and a5.3 cells at Stage 14 and Stage 20, respectively. White arrowheads indicate cleavage furrows. (E) Summary of the difference in cell cycle between a5.3 and A5.1 cell. (F) The cell volume of a5.3 cells in injected embryos at Stage 10. (G) The total surface area of a5.3 cells in injected embryos at Stage 10. (H) The cell volume of A5.1 cells in injected embryos at Stage 10. (I) The total surface area of A5.1 cells in injected embryos at Stage 10. (J) Representative images showing the cell long axis at stage 10. Top 50 long axes are shown using yellow (longest) to red (shortest) circles. Mesendoderm cells are outlined with white dots. Vegetal view. Anterior to the top, medial to the left. Rose diagrams showing the distribution of the longest axes relative to the medial plane of the embryo. (K) Orientation of mitotic spindles at Stage 14 in injected embryos. Embryos were stained with anti-α/ß-tubulin antibodies to visualize the mitotic spindle. Representative image of the cells are shown along with the cell long axes. (L) Rose diagrams showing the distribution of the longest axes relative to the medial plane of the embryo shown in Figure S4K. Angle of the spindle in the representative embryo is shown in orange. (M) The length of the mitotic spindle measured in 3D space. (N) Relationship between the angle between the spindle and the A5.1/A5.1 contact area to the angle between the spindle and the medial plane. (O) Relationship between the position of the furrow to the angle between the spindle and the medial plane. (P) Furrow position in response to the spindle angles. Red and blue planes correspond to the case of wee1 injected and control, respectively. Surfaces represent the model prediction using the best AIC model. C: Control RNA; TR: T19-Rho RNA. Scale bar = 10 µm (A, B, D), 50 µm (C, K). Numbers in the panels indicate the proportion of cells that are represented by the image. Scale bar = 50 µm. Error bars indicate one standard deviation.

**Supplemental Figure 5 Supplementary figure to Figure 5**

(A) The shape of the A5.1 cell when 20 pg of RNA encoding Q63L-Rho was injected into a5.3 cells. Note the excessive deformation of the cell toward the a5.3 cell. (B) Representative images showing the cell long axis at stage 10. Top 50 long axes are shown using yellow (longest) to red (shortest) circles. Mesendoderm cells are outlined with white dots. Vegetal view. Anterior to the top, medial to the left. (C) Rose diagrams showing the distribution of the longest axes relative to the medial plane of the embryo. (D) The cell volume of a5.3 cells in injected embryos at Stage 10. (E) The total surface area of a5.3-cells in injected embryos at Stage 10. (F) The cell volume of A5.1 cells in injected embryos at Stage 10. (G) The total surface area of A5.1 cells in injected embryos at Stage 10. (H) Orientation of mitotic spindles at Stage 14 in injected embryos. Embryos were stained with anti-α/ß-tubulin antibodies to visualize the mitotic spindle. Representative image of the cells are shown along with the cell long axes. Rose diagrams showing the distribution of the longest axes in the cells shown in Figure S5J. Angle of the spindle in the representative embryo is shown in orange. (I) Angle between the spindle and the medial plane in all the injected embryos. (J) Lengths of mitotic spindles measured in 3D space. (K) The position of the nucleus at Stage 7. Numbers in the panels indicate the proportion of cells that are represented by the image. Scale bar = 50 µm. Error bars indicate one standard deviation.

**Supplemental Movie 1: Movement of centrosomes and the nucleus**

(A) Formation of the cleavage furrow in the B4.2 cell. Embryos that displayed the same shape were selected as 0 min embryos under a stereomicroscope. (B) Mesendoderm cell (A5.1) and neighbor cell (a5.3) at stage 0, 1, and 2. (C) Cells at stage 3, 4, and 5. (D) Stage 6, 7, and 8 cells. (E) Stage 9, 10, and 11. (F) Stage 12, 13, and 14. (G) Closeup view of the centrosomes and the nucleus from stage 5, 6a (first 5 min), 6b (last 5 min) and 7. Nuclei were manually annotated yellow. Embryos were immunostained using anti-α/β- and γ-tubulin antibodies to visualize microtubules and centrosomes. The nucleus was manually annotated in red.

**Supplemental Movies 2: Position of the ME-plane during the 16-cell stage**

Time-lapse images of a representative 16-cell stage embryo. Embryos were stained using MitoTracker. Top row: image at the focal plane that includes the eCen. Bottom row: image at the focal plane that includes the nucleus. Left column: MitoTracker. Right column: overlay with bright view. Vegetal view, anterior to the top.

**Supplemental Movie 3: Position of the carbon label placed in the vicinity of the eCen**

3D movie showing the position of the carbon label in representative fixed embryos. Embryos were labeled at stage 7 and portion of the embryos were immediately fixed and stained using anti-α/β- and antibody to visualize the centrosome. Remainder of the embryos were cultured until stage 14 and fixed. The carbon labels were manually annotated red. To confirm the position of the carbon label, see Figure S1J. Animal pole to the top.

## Acknowledgements

Authors are grateful to fishermen in Aomori and Karakuwa for collecting adult Halocythia during the harsh conditions succeeding the 2011 earthquake and COVID-19, (ONO-SAN, CHIBA-SAN, ARIGATOU GOZAIMASHITA). NT would like to thank Dr. Kimiko Fukuda and members of the developmental biology laboratory, Tokyo Metropolitan University for helpful discussions and support in maintaining the laboratory. Authors would like to thank Dr. Tatsufumi Okamoto for critical reading of the draft and critical comments. NT would like to thank Dr. Jun-ichiro Suzuki of the plant ecology laboratory Tokyo Metropolitan University for kind and constructive advice on statistical analysis. NT would like to thank Professor Takahiro Asakage and his team in Tokyo Medical and Dental University for the operation they performed on NT’s thyroid. This study was supported by Japan Society for the Promotion of Science (25711016, 16H06169, 19K06694).

## Author contribution

Conceptualization, Data curation, Formal analysis, Investigation, Methodology, Software, Visualization, Writing: NT, YT; Funding acquisition, Resources, Project administration: NT.

