## Supplementary figures and images for "Cell cycle difference creates cortical tension difference that separates germ layer fates"

### Figure S1

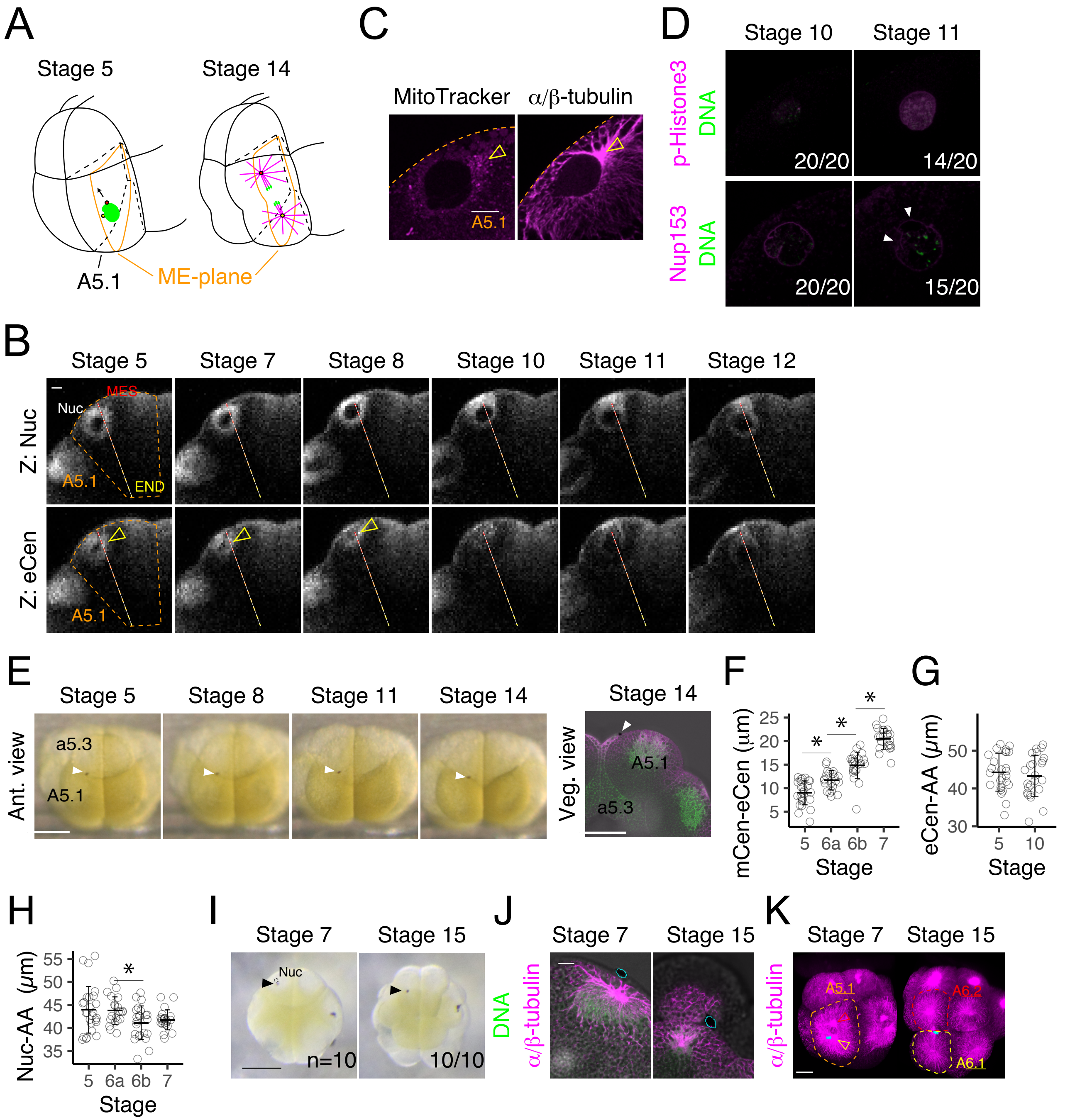

### Figure S2

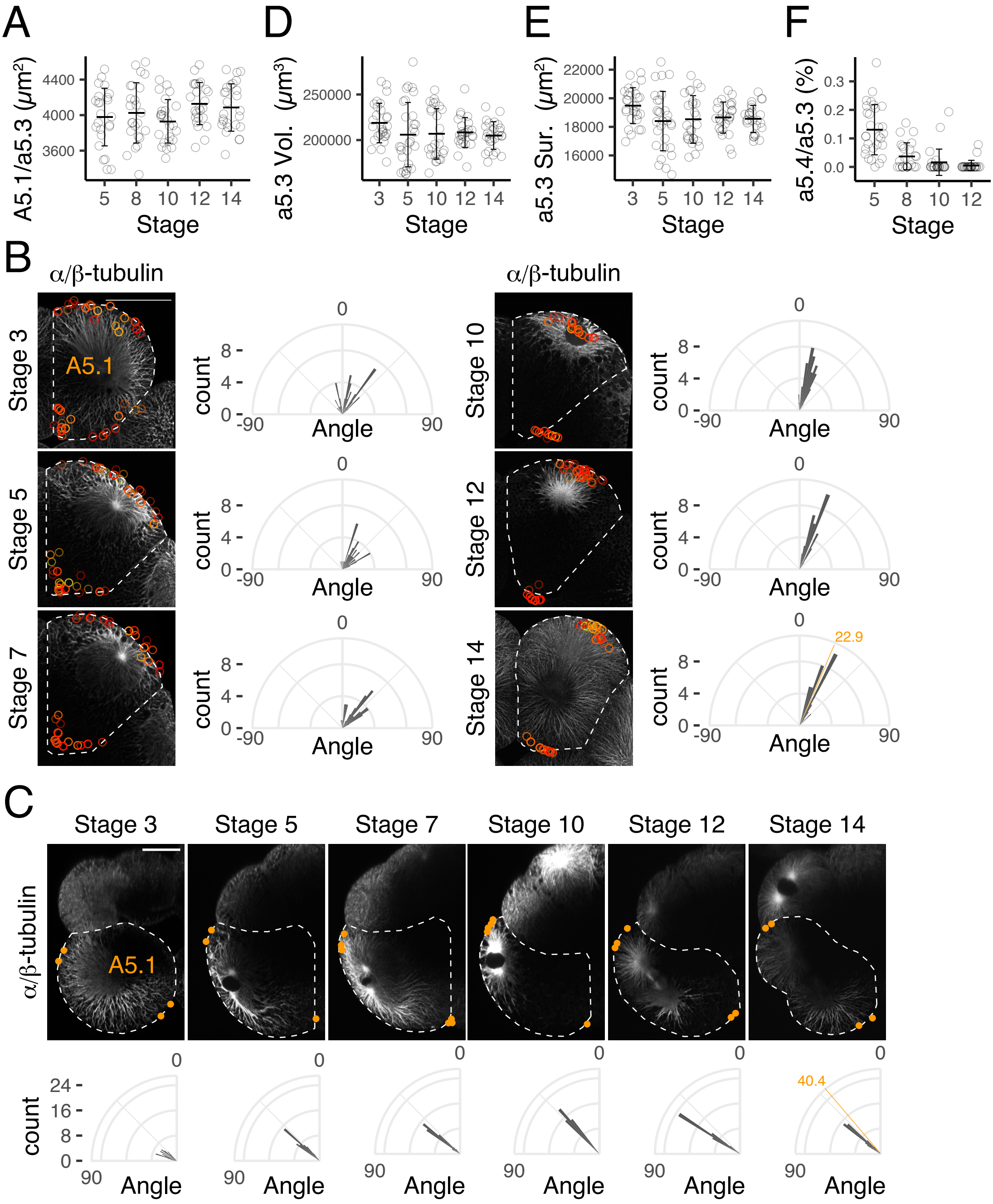

Figure S2

### Figure S3

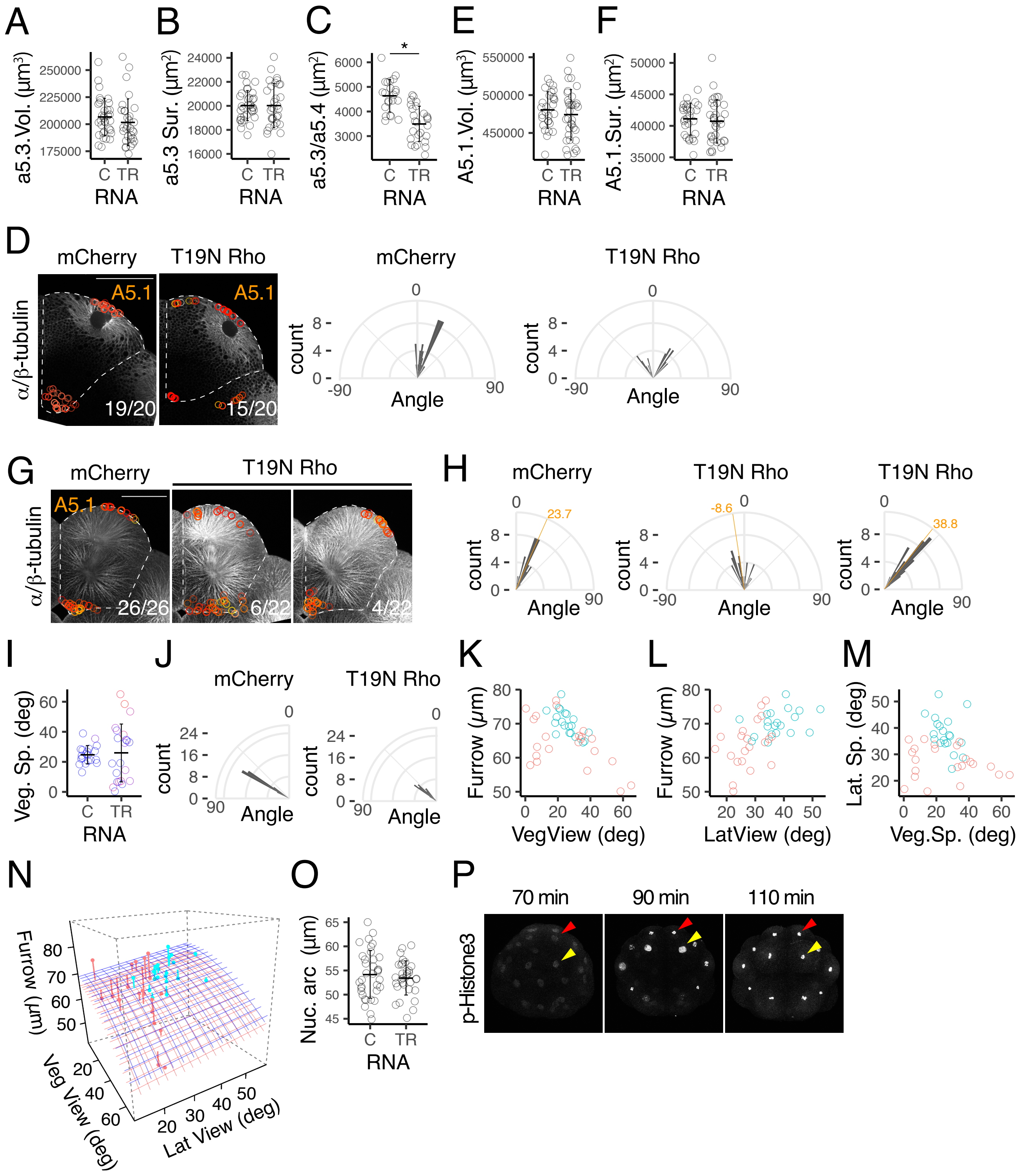

### Figure S4

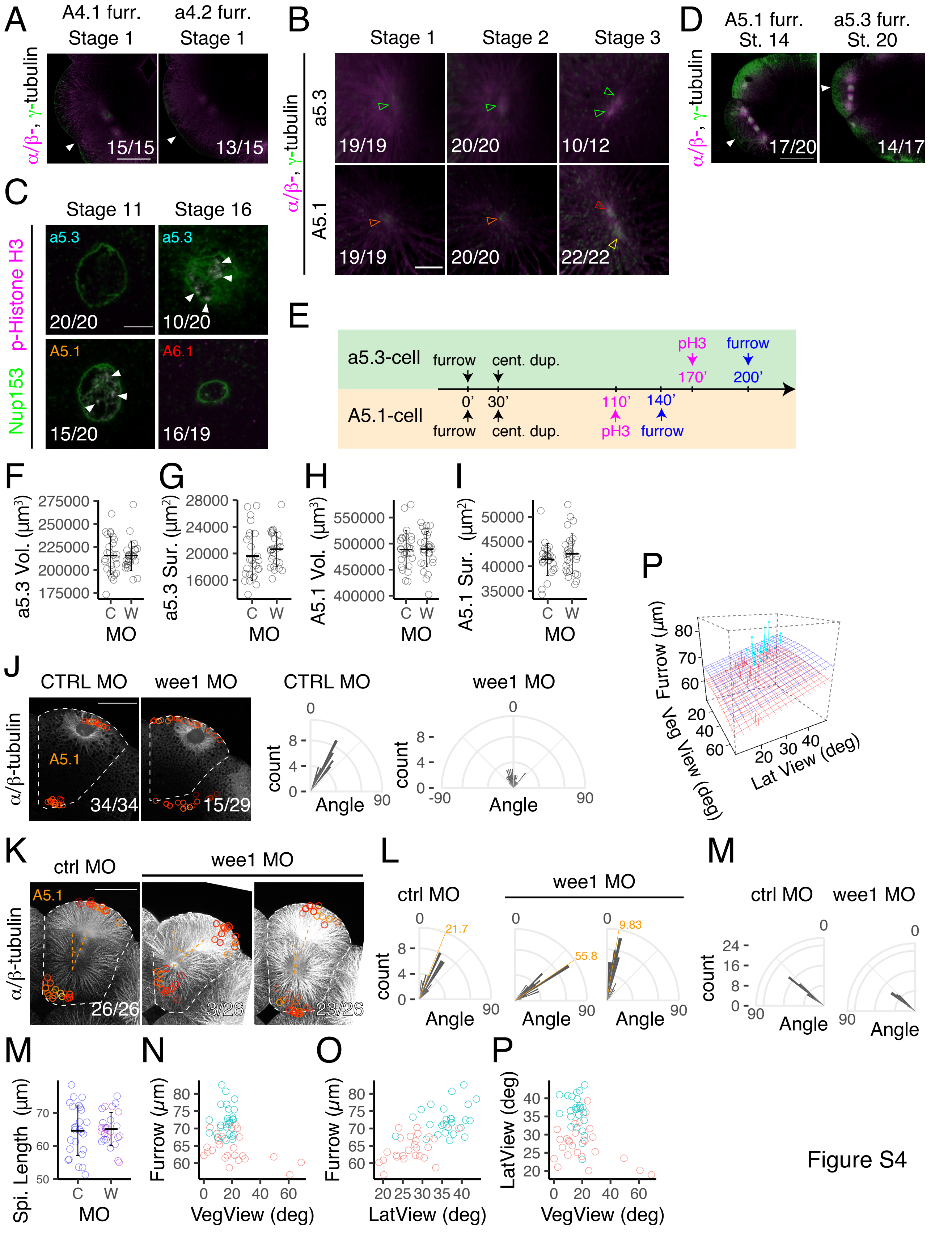

### Figure S5

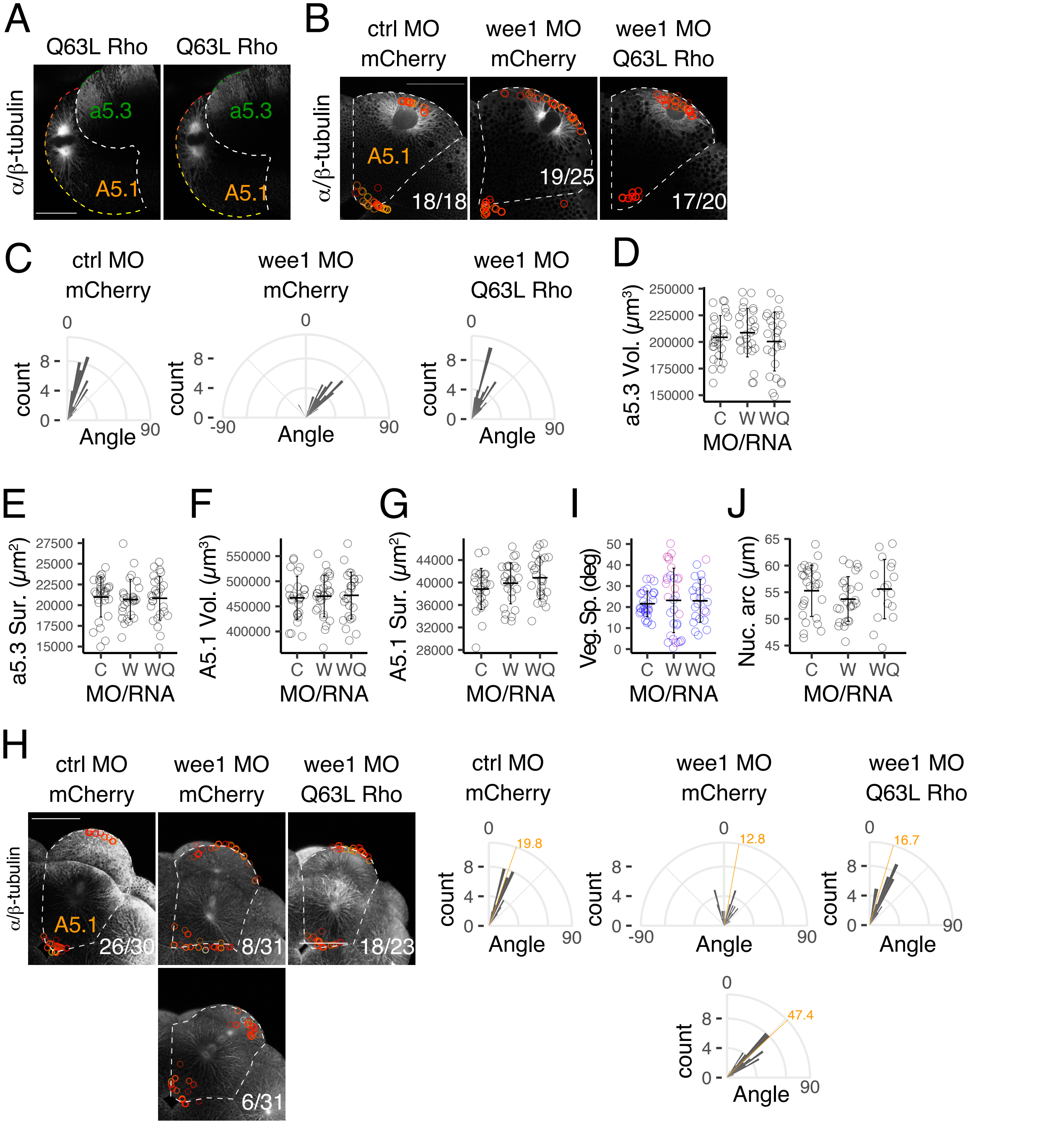
